# 16S rRNA amplicon sequencing for epidemiological surveys of bacteria in wildlife: the importance of cleaning post-sequencing data before estimating positivity, prevalence and co-infection

**DOI:** 10.1101/039826

**Authors:** Maxime Galan, Maria Razzauti, Emilie Bard, Maria Bernard, Carine Brouat, Nathalie Charbonnel, Alexandre Dehne-Garcia, Anne Loiseau, Caroline Tatard, Lucie Tamisier, Muriel Vayssier-Taussat, Helene Vignes, Jean François Cosson

## Abstract

Human impact on natural habitats is increasing the complexity of human-wildlife interfaces and leading to the emergence of infectious diseases worldwide. Highly successful synanthropic wildlife species, such as rodents, will undoubtedly play an increasingly important role in transmitting zoonotic diseases. We investigated the potential for recent developments in 16S rRNA amplicon sequencing to facilitate the multiplexing of large numbers of samples needed to improve our understanding of the risk of zoonotic disease transmission posed by urban rodents in West Africa. In addition to listing pathogenic bacteria in wild populations, as in other high-throughput sequencing (HTS) studies, our approach can estimate essential parameters for studies of zoonotic risk, such as prevalence and patterns of coinfection within individual hosts. However, the estimation of these parameters requires cleaning of the raw data to mitigate the biases generated by HTS methods. We present here an extensive review of these biases and of their consequences, and we propose a comprehensive trimming strategy for managing these biases. We demonstrated the application of this strategy using 711 commensal rodents collected from 24 villages in Senegal, including 208 *Mus musculus domesticus*, 189 *Rattus rattus*, 93 *Mastomys natalensis* and 221 *Mastomys erythroleucus.* Seven major genera of pathogenic bacteria were detected in their spleens: *Borrelia, Bartonella, Mycoplasma, Ehrlichia, Rickettsia, Streptobacillus* and *Orientia*. The last five of these genera have never before been detected in West African rodents. Bacterial prevalence ranged from 0% to 90% of individuals per site, depending on the bacterial taxon, rodent species and site considered, and 26% of rodents displayed coinfection. The 16S rRNA amplicon sequencing strategy presented here has the advantage over other molecular surveillance tools of dealing with a large spectrum of bacterial pathogens without requiring assumptions about their presence in the samples. This approach is therefore particularly suitable for continuous pathogen surveillance in the context of disease monitoring programs.

## Importance

Several recent public health crises have shown that the surveillance of zoonotic agents in wildlife is important to prevent pandemic risks. High-throughput sequencing (HTS) technologies are potentially useful for this surveillance, but rigorous experimental processes are required for the use of these effective tools in such epidemiological contexts. In particular, HTS introduces biases into the raw dataset that might lead to incorrect interpretations. We describe here a procedure for cleaning data before estimating reliable biological parameters, such as positivity, prevalence and coinfection, with 16S rRNA amplicon sequencing on the Illumina MiSeq platform. This procedure, applied to 711 rodents collected in West Africa, detected several zoonotic bacteria, including some at high prevalence despite never before having been reported for West Africa. In the future, this approach could be adapted for the monitoring of other microbes such as protists, fungi, and even viruses.

## Introduction

Pathogen monitoring in wildlife is a key method for preventing the emergence of infectious diseases in humans and domestic animals. More than half the pathogens causing disease in humans originate from animal species [1]. The early identification of zoonotic agents in animal populations is therefore of considerable interest for human health. Wildlife species may also act as a reservoir for pathogens capable of infecting livestock, with significant economic consequences [2]. The monitoring of emerging diseases in natural populations is also important for preserving biodiversity, because pathogens carried by invasive species may cause the decline of endemic species [3]. There is, therefore, a need to develop screening tools for identifying a broad range of pathogens in samples consisting of large numbers of individual hosts or vectors.

High-throughput sequencing (HTS) approaches require no prior assumptions about the bacterial communities present in samples of diverse nature, including non-cultivable bacteria. Such HTS microbial identification approaches are based on the sequencing of all (WGS: whole-genome sequencing) or some (RNAseq or 16S rRNA amplicon sequencing) of the bacterial DNA or RNA in a sample, followed by comparison to a reference sequence database [4]. HTS has made major contributions to the generation of comprehensive inventories of the bacteria, including pathogens, present in humans [5]. Such approaches are now being extended to the characterization of bacteria in wildlife [6–13]. However, improvements in the estimation of infection risks will require more than just the detection of bacterial pathogens. Indeed, we will also need to estimate the prevalence of these pathogens by host taxon and/or environmental features, together with coinfection rates [14,15] and pathogen interactions [16,17].

Razzauti *et al*. [8] recently used 16S rRNA amplicon sequencing with the dual-index sequencing strategy of Kozich *et al*. [18] to detect bacterial pathogens in very large numbers of rodent samples (up to several hundred samples in a single run) on the Illumina MiSeq sequencing platform. The 16S rRNA amplicon sequencing technique is based on the amplification of small fragments of one or two hypervariable regions of the 16S rRNA gene. The sequences of these fragments are then obtained and compared with reference sequences in curated databases for taxonomic identification [4,19]. Multiplexed approaches of this kind include short indices (or tags) linked to the PCR products and specific to a given sample. This makes it possible to assign the sequences generated by the HTS run to a particular sample following bioinformatic analysis of the dataset [18]. Razzauti *et al*. [8] demonstrated the considerable potential of this approach for determining the prevalence of bacteria within populations and for analyzing bacterial interactions within hosts and vectors, based on the accurate characterization of bacterial diversity within each individual samples it provides. However, various sources of error during the generation and processing of HTS data [20] may make it difficult to determine which samples are really positive or negative for a given bacterium. The detection of one or a few sequences assigned to a given taxon in a sample does not necessarily mean that the bacterium is actually present in that sample. We carried out an extensive literature review, from which we identified several potential sources of error involving all stages of a 16S rRNA amplicon sequencing experiment — from the collection of samples to the bioinformatic analysis — that might lead to false-negative or false-positive screening results (Table 1, [18,19,21–40]). These error sources have now been documented, and recent initiatives have called for the promotion of open sharing of standard operating procedures and best practices in microbiome research [41]. However, no experimental designs minimizing the impact of these sources of error on HTS data interpretation have yet been reported.

We describe here a rigorous experimental design for the direct estimation of biases from the data produced by 16S rRNA amplicon sequencing. We used these bias estimates to control and filter out potential false-positive and false-negative samples during screening for bacterial pathogens. We applied this strategy to 711 commensal rodents collected from 24 villages in Senegal, Western Africa: 208 *Mus musculus domesticus*, 189 *Rattus rattus*, 93 *Mastomys natalensis* and 221 *Mastomys erythroleucus*. Pathogenic bacteria associated with the rodents were analysed using a protocol based on Illumina MiSeq sequencing of the V4 hypervariable region of the 16S rRNA gene [18]. We considered the common pitfalls listed in Table 1 during the various stages of the experiment (see details in the workflow procedure, Figure 1). Biases in assessments of the presence or absence of bacteria in rodents were estimated directly from the dataset, by including and analysing negative controls (NC) and positive controls (PC) at various stages of the experiment (see Box 1), and systematically using sample replicates. This strategy delivers realistic and reliable estimates of bacterial prevalence in wildlife populations, and could be used to analyse the co-occurrence of different bacterial species within individuals.

**Table 1.**
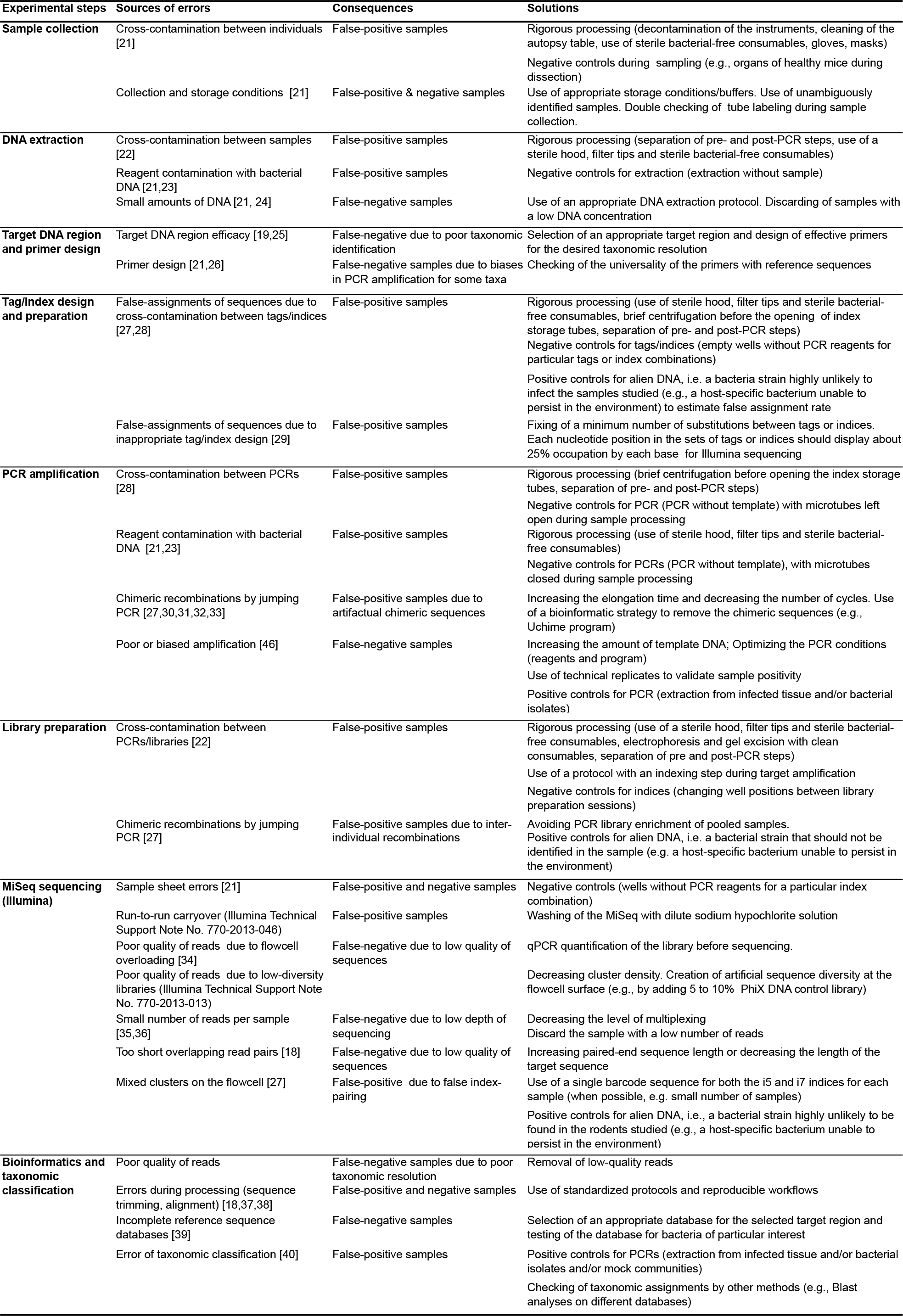
**Sources of bias during the experimental and bioinformatic steps of 16S rRNA amplicon sequencing.** Consequences for data interpretation and solutions for mitigating these biases.

**Figure 1.**
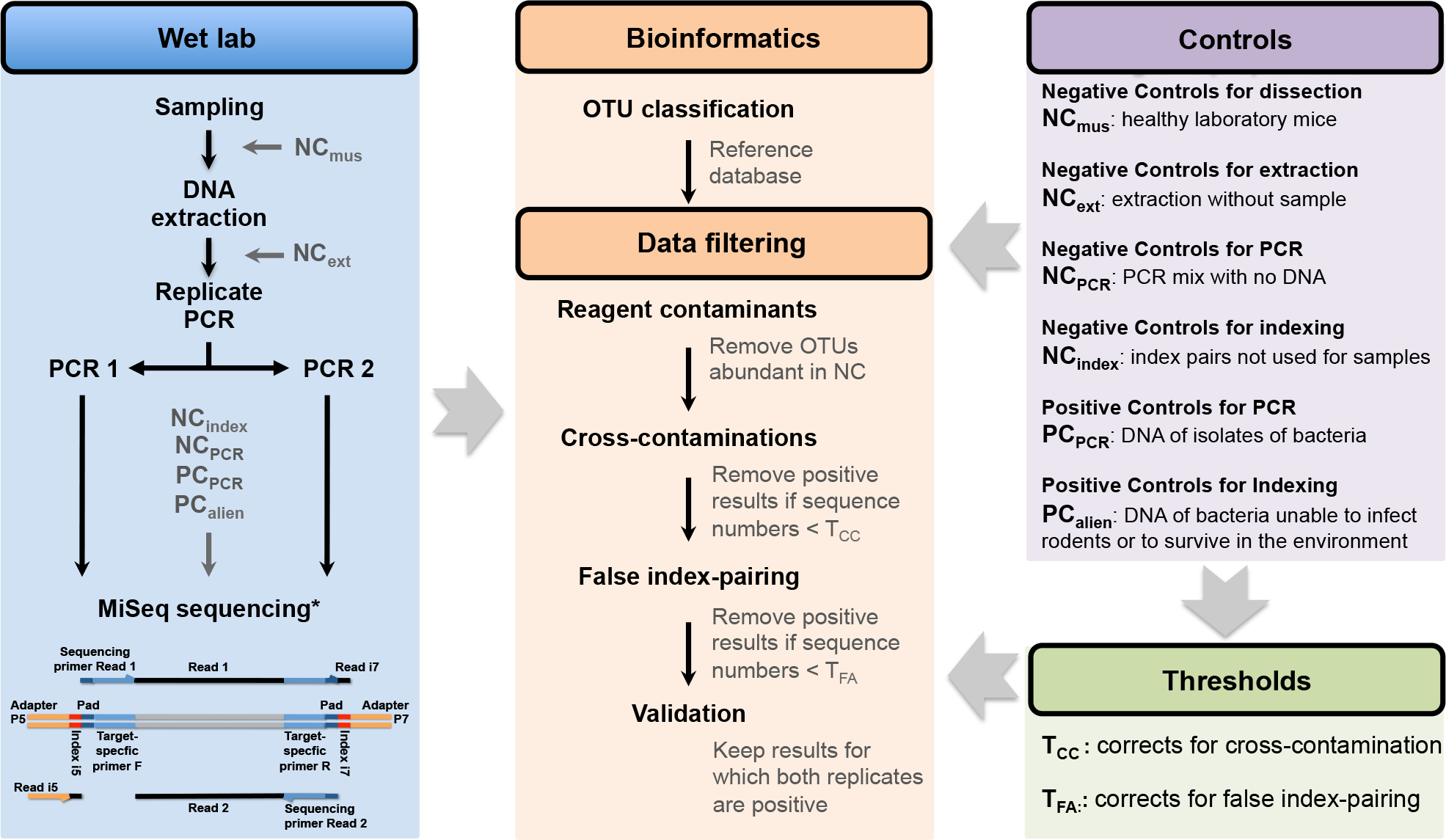
**Workflow of the wet laboratory, bioinformatics and data filtering procedures in the process of data filtering for 16S rRNA amplicon sequencing.** Reagent contaminants were detected by analyzing the sequences in the NC_ext_ and NC_PCR_ controls. Sequence number threshold for correcting for cross-contamination (T_CC_) are OTU-and run-dependent, and were estimated by analyzing the sequences in the NC_mus_, NC_ext_, NC_PCR_ and PC_index_ controls. Sequence number threshold for correcting for false index-pairing (T_FA_) values are OTU-and run-dependent, and were estimated by analyzing the sequences in the NC_index_ and PC_allen_ controls. A result was considered positive if the number of sequences was > T_CC_ and > T_FA_. Samples were considered positive if a positive result was obtained for both PCR replicates. *see Kozich et al 2013 for details on the sequencing.

## Results & Discussion

***Raw sequencing results.*** The sequencing of 1569 PCR products in two MiSeq runs generated a total of 23,698,561 raw paired-end sequence reads (251-bp) of the V4 region of the 16S rRNA gene. Because we made PCR replicates for each rodent sample, and because we included several controls in each sequencing run, we have more PCR products (N=1569) than rodent samples (N=711) (see summary in Table S1 and complete information by sample and run in Table S2). Overall, 99% of PCRs generated more than 3,000 raw reads (mean: 11,908 reads; standard deviation: 6,062). The raw sequence files are available in FASTQ format in the Dryad Digital Repository http://dx.doi.org/10.5061/dryad.m3p7d [42].

Using mothurv1.34 [43] and the MiSeq standard operating procedure (http://www.mothur.org/wiki/MiSeq_SOP), we removed 20.1% of paired-end reads because they were misassembled, 1.5% of sequences because they were misaligned, 2.6% because they were chimeric and 0.2% because they were non-bacterial. The remaining reads were grouped into operational taxonomic units (OTUs) with a divergence threshold of 3%. Bioinformatics analysis identified 13,296 OTUs, corresponding to a total of 7,960,533 sequences in run 1 and 6,687,060 sequences in run 2.

#### Box 1. Guideline for experimental controls to include within high-throughput amplicon sequencing experiments to mitigate false positive results

Recent research has highlighted different biases occurring at different steps of high-throughput sequencing. These biases can be estimated directly from the data by including several controls together with samples in the experiment. We detail below these different controls as well as the rationale for their use.

**Negative Controls for sample collection.** When possible we advise to include axenic samples during sample collection. The number of sequences observed in these controls are used to estimate cross-contamination rates during sample collection. In our study we used spleens from healthy laboratory mice (**NC**_mus_), free from rodent pathogens, which were manipulated together with wild samples during the dissections in the field.

**Negative Controls for DNA extraction (NC_ext_).** DNA extractions performed without the addition of sample tissue (blanks), which are processed together with the other samples. We advise performing at least one extraction blank for each extraction experiment, although more is better. The numbers of sequences observed in these controls are used to estimate and filter the cross-contaminations during the DNA extractions and to detect for DNA bacterial contaminants in the extraction kit reagents.

**Negative Controls for PCR (NC_pcr_).** PCR reactions without any DNA extract included (blank), which are processed together with the other samples. We advise performing at least one PCR blank per PCR microplate, although more is better. The numbers of sequences observed in these controls are used to estimate and filter the cross-contaminations during the PCR preparation and to detect DNA bacterial contaminants in the PCR reagents.

**Negative Controls for indexing (NC_index_).** Combinations of barcodes that are not used to identify samples in the sequencing run, but that are searched for during the bioinformatic demultiplexing. In practice, they correspond to empty PCR wells (without reagent and without index). The numbers of sequences recovered for these particular index combinations are used to estimate and filter the cross-contaminations between indexed PCR primers during primer handling or PCR preparation, and to identify errors in the Illumina sample sheet.

**Positive Controls for PCR (PC_PCR_).** PCR reactions with DNA of known taxa isolates, which are processed together with the other samples. The sequences obtained for these controls are used to verify the taxonomic assignment and to estimate and filter cross-contaminations.

**Positive Controls for Indexing (PC_alien_).** PCR reactions with DNA of taxa isolates that are known to be absent in the samples. They are handled separately from the samples to avoid cross-contaminations with the samples during the wet lab procedures (DNA extractions and PCRs). Sequences from PC_aNen_ found in the samples are used to calculate the rate of sample misidentification due to false index-pairing (see text and Kircher et al [27] for details concerning this phenomenon).

In practice, (PC_PCR_) and (PC_alien_) could be the same and we advice to use taxa that are phylogenetically distant from the taxa we look for, in order to avoid potential confusion between sequences from alien controls and sequences from the samples.

***Taxonomic assignment of sequences.*** We used the Bayesian classifier (bootstrap cutoff = 80%) implemented in mothurwith the Silva SSU Ref database v119 [43] as a reference, for the taxonomic assignment of OTUs. The 50 most abundant OTUs accounted for 89% (min: 15,284 sequences; max: 2,206,731 sequences) of the total sequence dataset (Table S3). The accuracy of taxonomic assignment (to genus level) was assessed with positive controls for PCR, corresponding to DNA extracts from laboratory isolates of *Bartonella taylorii, Borrelia burgdorferi* and *Mycoplasma mycoides* (PC_Bartonella_t_, PC_Borrelia_b_ and PC_Mycoplasma_m_, respectively), which were correctly assigned to a single OTU corresponding to the appropriate reference sequences (Table 2). Note that the sequences of PC_Mycoplasma_m_ were assigned to Entomoplasmataceae rather than Mycoplasmataceae because of a frequent taxonomic error reflected in most databases, including Silva [45]. This problem might also affect other taxa. We therefore recommend systematically carrying out a blast analysis against the sequences of taxa of interest in GenBank to confirm the taxonomic assignment obtained with the 16S databases. Finally, we assumed that the small number of sequences per sample might limit the completeness of bacterial detection [36]. For this reason, we discarded seven rodent samples (2 *M. erythroleucus* and 5 *M. domesticus)* yielding fewer than 500 sequences for at least one of the two PCR replicates (1% of the samples).

**Table 2.**
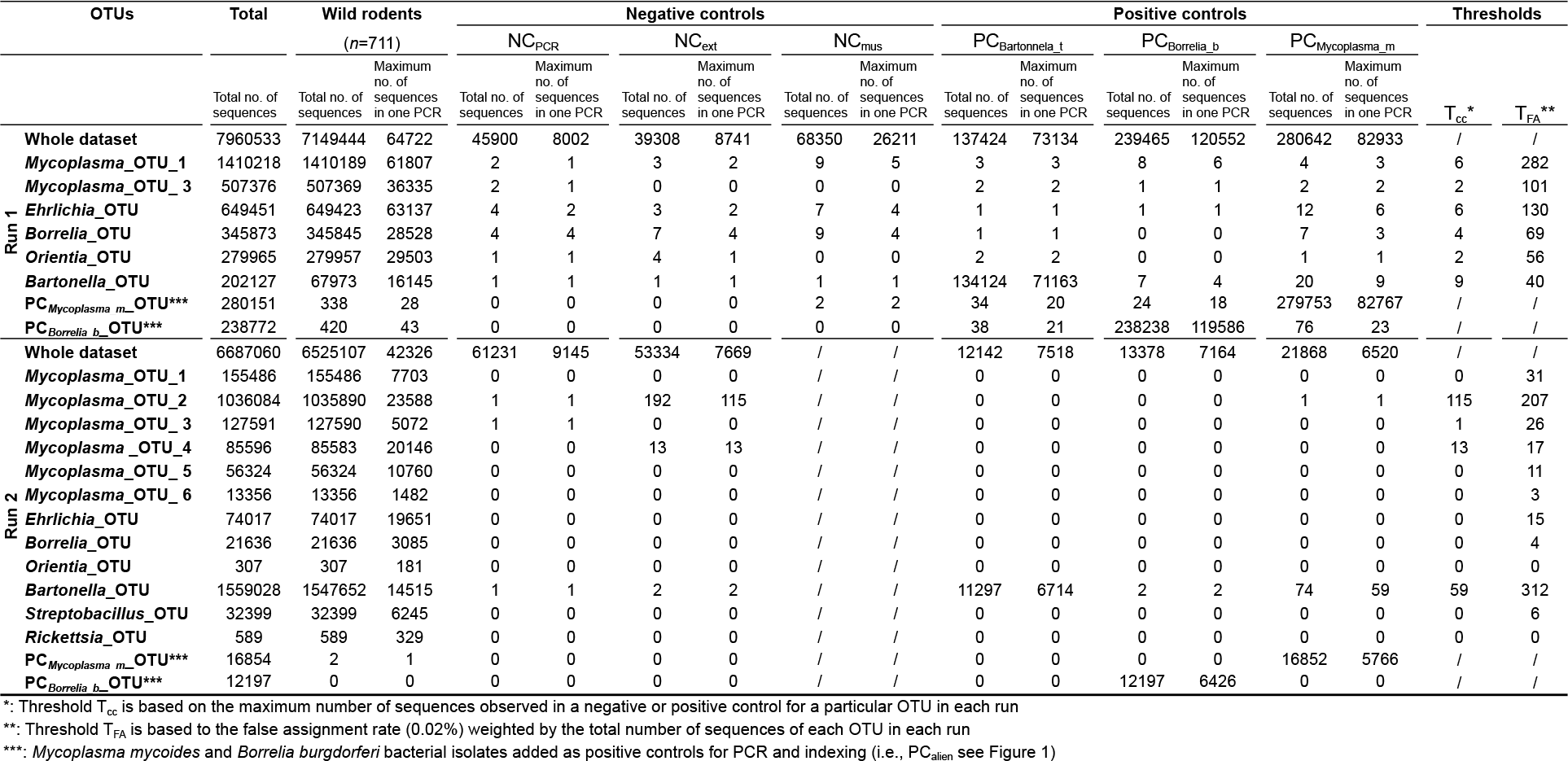
**Number of sequences for 12 pathogenic OTUs observed in wild rodents, negative controls, and positive controls, together with T_CC_ and T_FA_ threshold values.** Data are given for the two MiSeq runs separately. NC_pCr_: negative controls for PCR; NC_ext_: negative controls for extraction; NC_mus_: negative controls for dissection; PC_Bartoneiia_t_: positive controls for PCR; PC_Borreiia_b_ and PC_My_co_p_is_m_a__m_: positive controls for PCR and positive controls for indexing; T_cc_ and T_FA_: thresholds for positivity for a particular bacterium according to bacterial OTU and MiSeq run (see also Figure 1).

***Filtering for reagent contaminants.*** 16S rRNA amplicon sequencing data may be affected by the contamination of reagents [23], We therefore filtered the data, using negative controls for extraction (NC_ext_), corresponding to extraction without the addition of a tissue sample, and negative controls for PCR (NC_PCR_), corresponding to PCR mixtures to which no DNA was added. We observed between 2,843 and 8,967 sequences in the NC_ext_ and between 5,100 and 9,145 sequences in the NC_PCR_. Based on their high number of reads in negative controls, we identified 13 contaminant genera, including *Pseudomonas, Acinetobacter, Herbaspirillum, Streptococcus, Pelomonas, Brevibacterium, Brachybacterium, Dietzia, Brevundimonas, Delftia, Comamonas, Corynebacterium*, and *Geodermatophilus*, some of them having been previously identified in other studies [23], These contaminants accounted for 29% of the sequences in the dataset (Figure 2). They also differed between MiSeq runs: *Pseudomonas, Pelomonas* and Herbaspirillum predominated in run 1, whereas *Brevibacterium, Brachybacterium* and *Dietzia* predominated in run 2 (Table S4, Figure S1). This difference probably reflects the use of two different PCR kits manufactured several months apart (Qiagen technical service, pers. com.). The majority of other contaminants, such as *Streptococcus*, most likely originated from the DNA extraction kits used, as they were detected in abundance in the negative controls for extraction (NC_ext_).

**Figure 2.**
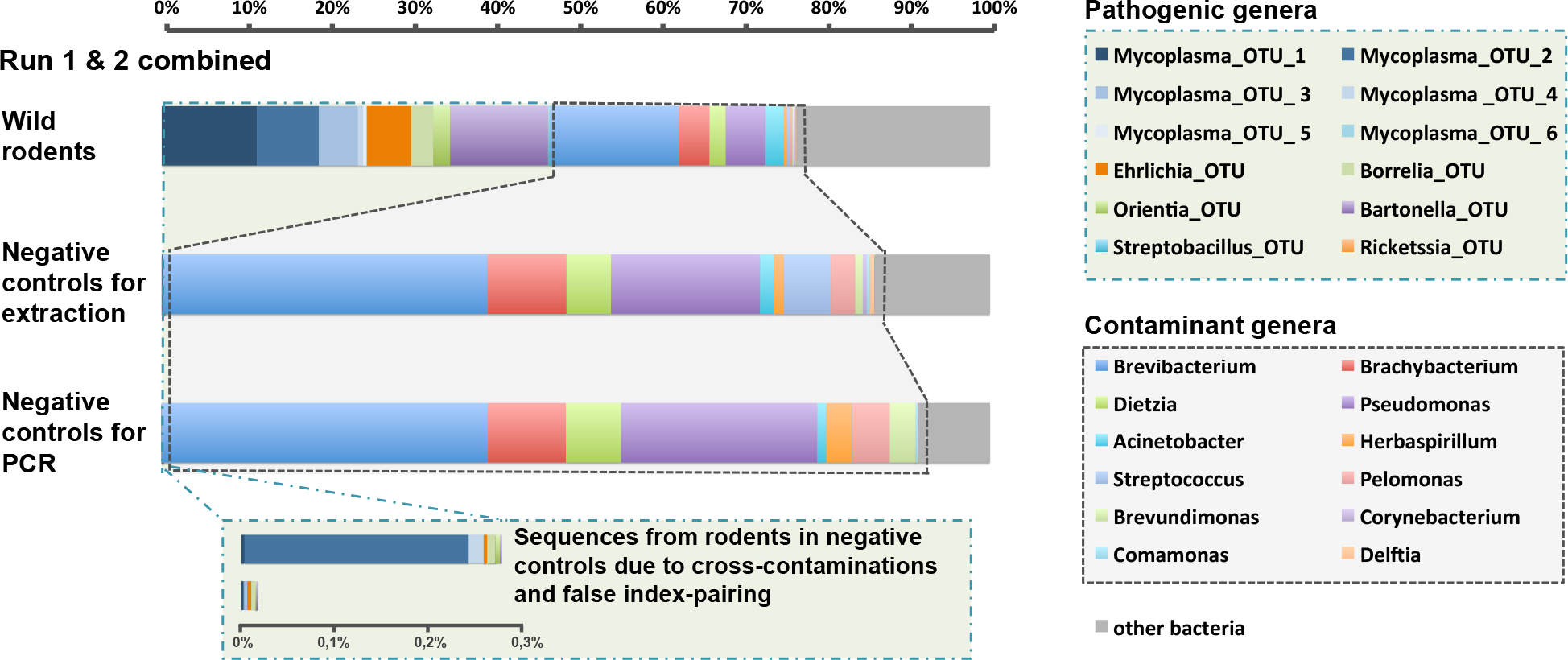
**Taxonomic assignment of the V4 16S rRNA sequences in wild rodents, and in negative controls for extraction and PCR.** The histograms show the percentage of sequences for the most abundant bacterial genera in the two MiSeq runs combined. Notice the presence of several bacterial genera in the controls, which were likely due to the inherent contamination of laboratory reagents by bacterial DNA (termed ‘contaminant genera’). These contaminant genera are also present (to a lesser extent) in the rodent samples. The inserts represent the proportion of sequences from rodent samples, which were incorrectly assigned to the controls. See Figure S1 for separate histograms for both MiSeq runs.

Genera identified as contaminants were then simply removed from the sample dataset. It is important to note, however, that the exclusion of these results does not rule out the possibility that our samples contained true rodent infections (at least for some of them like *Streptococcus* which contains both saprophytic and pathogenic species). However, as mentioned by Razzauti *et al*. [8] distinguishing between those two possibilities seems difficult, if not impossible. Faced with this lack of certainty, it is most prudent to simply remove these taxa from the sample dataset. These results highlight the importance of carrying out systematic negative controls to filter the taxa concerned in order to prevent inappropriate data interpretation, particularly for the *Streptococcus* genus, which contains a number of important pathogenic species. The use of DNA-free reagents would improve the quality of sequencing data and likely increase the depth of sequencing of the samples.

After filtering for the above reagent contaminants, 12 OTUs, belonging to 7 genera for which at least one species or one strain is known to be pathogenic in mammals (therefore referenced as “pathogenic genera”), accounted for 66% of the sequences identified in wild rodent samples for both MiSeq runs combined (Figure 2). These genera are *Bartonella, Borrelia, Ehrlichia, Mycoplasma, Orientia, Rickettsia* and *Streptobacillus*. Six different OTUs were obtained for *Mycoplasma (Mycoplasma_OTU_1* to *Mycoplasma_OTU_6)*, and one OTU each for the other genera (Table 2). Finally, the precise significance of the remaining 34% of sequences was undetermined, potentially corresponding to commensal bacteria (Bacteroidales, Bacteroides, Enterobacteriaceae, Helicobacter, Lactobacillus), unknown pathogens, undetected contaminants, or undetected sequencing errors.

***Filtering for false-positive results.*** Mothur analysis produced a table of abundance, giving the number of sequences for each OTU in each PCR product (data available in the Dryad Digital Repository http://dx.doi.org/10.5061/dryad.m3p7d [42]. The multiple biases during experimental steps and data processing listed in Table 1 made it impossible to infer prevalence and co-occurrence directly from the table of sequence presence/absence in the PCR products. We suggest filtering the data with estimates of the different biases calculated from the multiple controls introduced during the process. This strategy involves calculating sequence number thresholds from our bias estimates. Two different thresholds were set for each of the 12 OTUs and two MiSeq runs. We then discarded positive results associated with sequence counts below the threshold (Figure 1).

***Threshold T_cc_: Filtering for cross-contamination.*** One source of false positives is cross-contamination between samples processed in parallel (Table 1). Negative controls for dissection (NC_mus_), consisting of the spleens of healthy laboratory mice manipulated during sessions of wild rodent dissection, and negative controls for extraction (NC_ext_) and PCR (NC_PCR_) were used, together with positive controls for PCR (PC_Bartonella_t_, PC_Borrelia_b_ and PC_Mycoplasma_m_), to estimate cross-contamination. For each sequencing run, we calculated the maximal number of sequences for the 12 pathogenic OTUs in the negative and positive controls. These numbers ranged from 0 to 115 sequences, depending on the OTU and the run considered (Table 2), and we used them to establish OTU-specific thresholds (T_cc_) for each run. For example, in Sequencing Run 2, the highest number of sequences in a control for *Mycoplasma_OTU_2* was 115 (in a NC_ext_). Therefore, we established the threshold value at 115 sequences for this OTU in sequencing Run 2. Thus, PCR products with less than 115 sequences for the *Mycoplasma_OTU_2* in sequencing Run 2 were considered as false-positive for this OTU. The use of these T_cc_ led to 0% to 69% of the positive results being discarded, corresponding to only 0% to 0.14% of the sequences, depending to the OTU considered (Figure 3, Table S5). A PCR product may be positive for several bacteria in cases of coinfection. In such cases, the use of a T_cc_ makes it possible to discard the positive result for one bacterium whilst retaining positive results for other bacteria.

***Threshold T_FA_: Filtering out incorrectly assigned sequences.*** Another source of false positives is the incorrect assignment of sequences to a PCR product (Table 1). This phenomenon may be due either to cross-contamination between indices during the experiment, or to the generation of mixed clusters during the sequencing [27].

First, the cross-contamination of indexes may happen during the preparation of indexed primer microplates. This cross-contamination was estimated using negative control index pairs (NC_index_) corresponding to particular index pairs not used to identify the samples. NC_index_ returned very few read numbers (1 to 12), suggesting that there was little or no cross-contamination between indices in our experiment.

Second, the occurrence of mixed clusters during the sequencing of multiplexed samples was reported by Kircher et al [27]. Mixed clusters on the Illumina flowcell surface are considered by Kircher et al [27] as the predominant source of error of sequence assignment to a PCR product. The impact of this phenomenon on our experiment was estimated using “alien” positive controls (PC_alien_) sequenced in parallel of the rodent samples: PC_Mycoplasma_m_, corresponding to the DNA of *Mycoplasma mycoides*, which cannot infect rodents, and PC_Borrelia_b_, containing the DNA of *Borrelia burgdorferi*, which is not present in Africa. Neither of these bacteria can survive in abiotic environments, so the presence of their sequences in African rodent PCR products indicates a sequence assignment error due to false index-pairing [27]. Using PC_Mycoplasma_m_, we obtained an estimate of the global false index-pairing rate of 0.14% (i.e. 398 of 280,151 sequences of the *Mycoplasma mycoides* OTU were assigned to samples other than PC_Mycoplasma_m_). Using PC_Borrelia_b_, we obtained an estimate of 0.22% (534 of 238,772 sequences of the *Borrelia burgdorferi* OTU were assigned to samples other than PC_Borrelia_b_). These values are very close to the estimate of 0.3% obtained by Kircher *et al*. [27]. Close examination of the distribution of misassigned sequences within the PCR 96-well microplates showed that all PCR products with misassigned sequences had one index in common with either PC_Mycoplasma_m_ or PC_Borrelia_b_ (Figure S2).

We then estimated the impact of false index-pairing for each PCR product by calculating the maximal number of sequences of “alien” bacteria assigned to PCR products other than the corresponding PC. These numbers varied from 28 to 43, depending on the positive control for run 1 (Table 2) — run 2 was discarded because of the low values of the numbers of sequences, which is likely due to the fact that DNAs of PC were diluted one hundred-fold in run 2 (Table S1). We then estimated a false-assignment rate for each PCR product (R_fa_), by dividing the above numbers by the total number of sequences from “alien” bacteria in Sequencing Run 1. R_fa_ was estimated for PC_MycoplaS_ma_m and PC_Borrelia_b_ separately. R_fa_ reached 0.010% and 0. 018% for PC_Mycoplasma_m_ and PC_Borrelia_b_, respectively. We adopted a conservative approach, by fixing the R_fa_ value to 0.020%. This number signifies that each PCR product may receive a maximum 0.020% of the total number of sequences of an OTU present in a run due to false index-pairing. Moreover, the number of misassigned sequences for a specific OTU into a PCR product should increase with the total number of sequences of the OTU in the MiSeq run. We therefore defined the second threshold (T_FA_) as the total number of sequences in the run for an OTU multiplied by R_fa_. TFA values varied with the abundance of each OTU in the sequencing run (Table 2). Because the abundance of each OTU varied from one sequencing run to the other, T_FA_ also varied according to the sequencing run. The use of the T_FA_ led to 0% to 87% of positive results being discarded. This corresponded to 0% to 0.71% of the sequences, depending on the OTU (Figure 3, Table S5).

**Figure 3.**
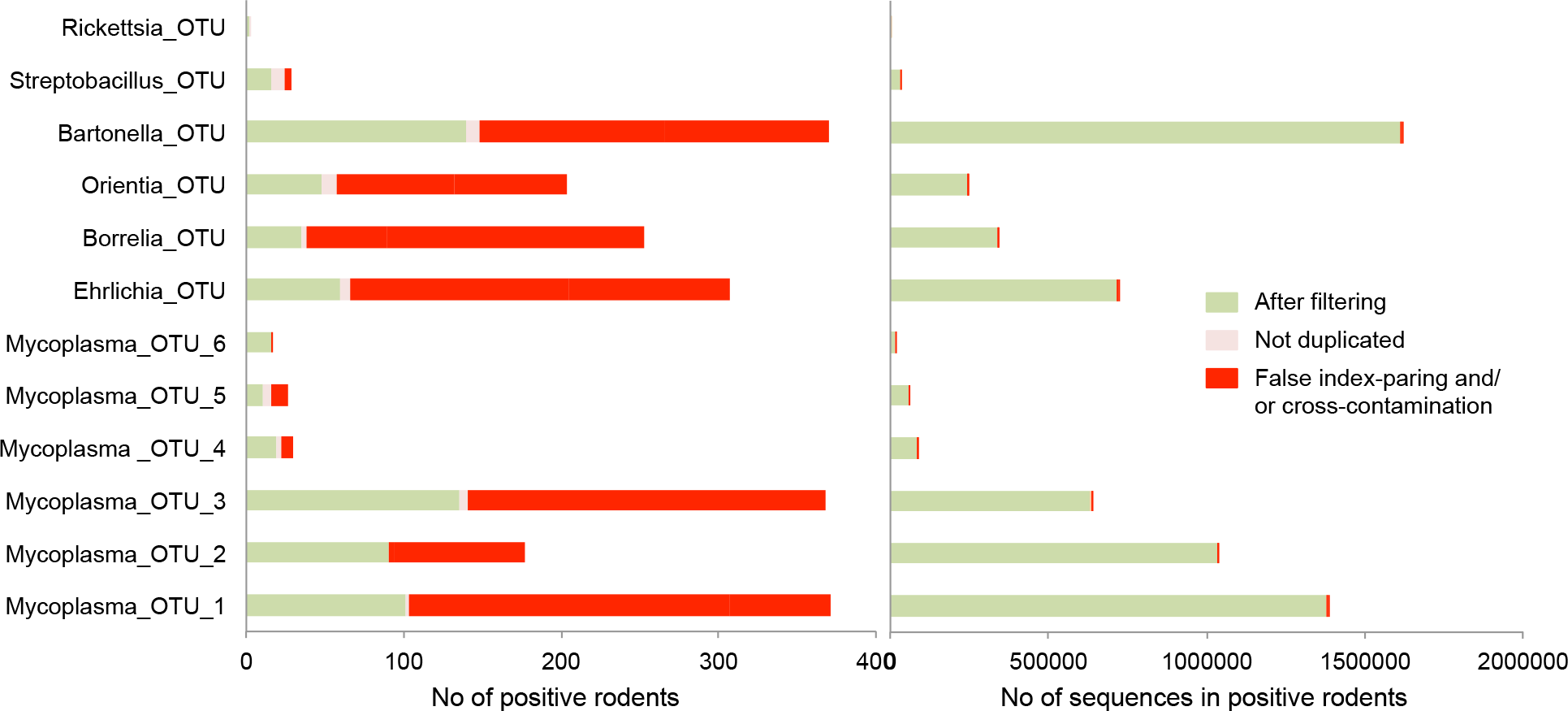
**Numbers of positive rodents, and of sequences in positive rodents, removed for each OTU at each step in data filtering.** These findings demonstrate that the positive rodents filtered out corresponded to only a very small number of sequences. (A) The histogram shows the number of positive rodents discarded because of likely cross-contamination, false index-pairing, and failure to replicate in both PCRs, as well as the positive results retained at the end of data filtering in green. (B) The histogram shows the number of sequences corresponding to the same class of positive rodents. Note that several positive results may be recorded for the same rodent in cases of co-infection

***Validation with PCR replicates.*** Random contamination may occur during the preparation of PCR 96-well microplates. These contaminants may affect some of the wells, but not those for the negative controls, leading to the generation of false-positive results. We thus adopted a conservative approach, in which we considered rodents to be positive for a given OTU only if both PCR replicates were considered positive after the filtering steps described above. The relevance of this strategy was supported by the strong correlation between the numbers of sequences for the two PCR replicates for each rodent (R^2^>0.90, Figure 4 and Figure S3). At this stage, 673 positive results for 419 rodents were validated for both replicates (note that a rodent may be positive for several bacteria, and may thus be counted several times), whereas only 52 positive results were discarded because the result for the other replicate was negative. At this final validation step, 0% to 60% of the positive results for a given OTU were discarded, corresponding to only 0% to 7.17% of the sequences (Figure 3, Table S5 and Table S6). Note that the number of replicates may be increased, as described in the strategy of Gomez-Dfaz *et al* [46].

**Figure 4.**
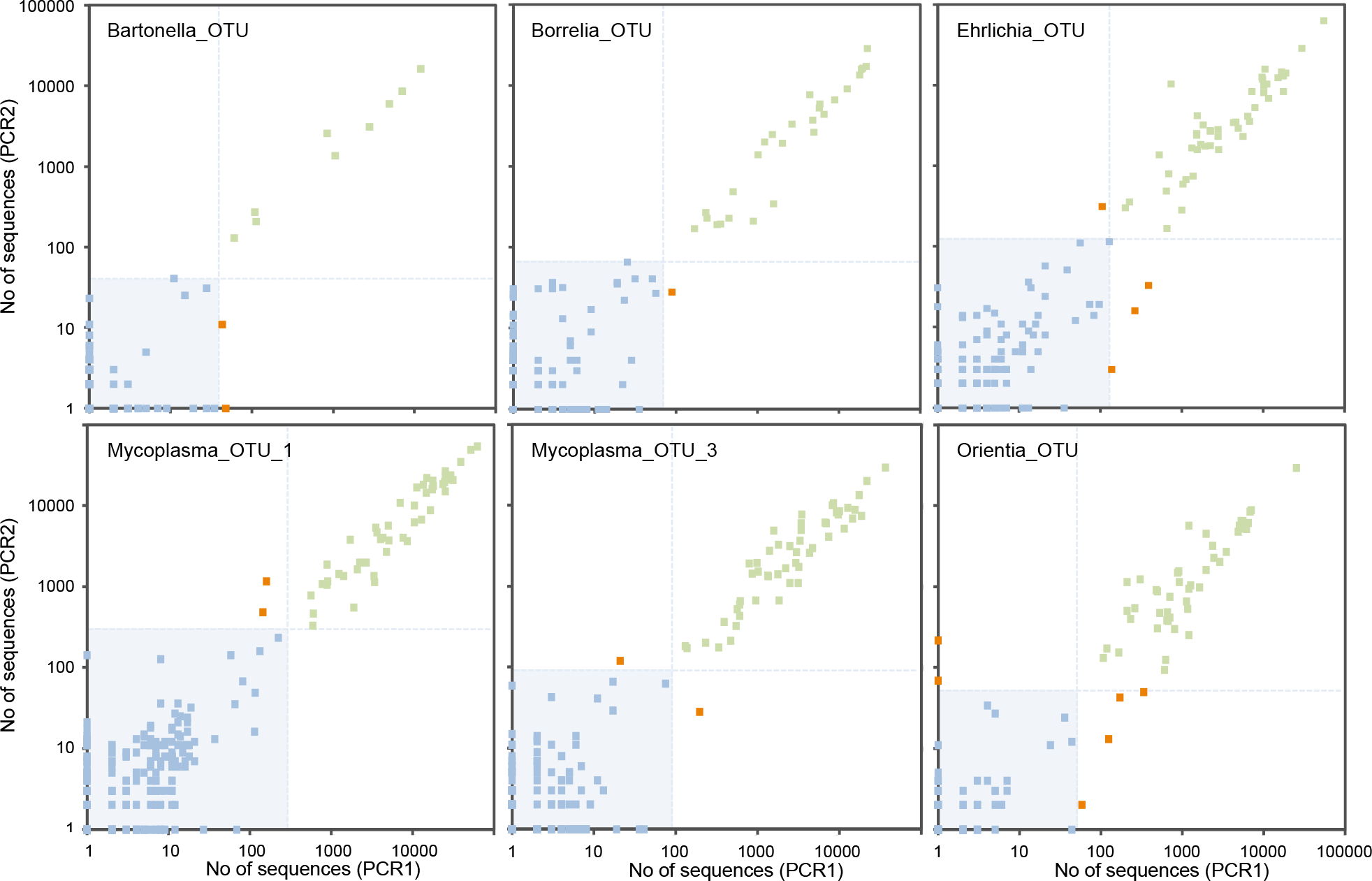
**Plots of the number of sequences (log (x+1) scale) from bacterial OTUs in both PCR replicates (PCR1 & PCR2) of the 348 wild rodents analyzed in the first MiSeq run.** Note that each rodent was tested with two replicate PCRs. Green points correspond to rodents with two positive results after filtering; red points correspond to rodents with one positive result and one negative result; and blue points correspond to rodents with two negative results. The light blue area and lines correspond to threshold values used for the data filtering: samples below the lines are filtered out. See Figure S3 for plots corresponding to the second MiSeq run.

***Post-filtering results.*** Finally, the proportion of rodents positive for a given OTU filtered out by the complete filtering approach varied from 6% to 86%, depending on the OTU, corresponding to only 1% of the total sequences (Figure 3). Indeed, our filtering strategy mostly excluded rodents with a small number of sequences for the OTU concerned. These rodents were considered to be false-positive.

***Refining bacterial taxonomic identification.*** We refined the taxonomic identification of the 12 bacterial OTUs through phylogenetic and blast analyses. We were able to identify the bacteria present down to genus level and, in some cases, we could even identify the most likely species (Table 3 and Figure S4). For instance, the sequences of the six *Mycoplasma* OTUs were consistent with three different species — *M. haemomuris* for OTU_1 and 3, *M. coccoides* for OTU_4, 5 and 6, and *M. species novo* [47] for OTU_2 — with high percentages of sequence identity (≥93%) and bootstrap values ≥80%. All three of these species belong to the Hemoplasma group, which is known to infect mice, rats and other mammals [48,49], and is thought to cause anemia in humans [50,51]. The *Borrelia* sequences grouped with three different species of the relapsing fever group (*crocidurae, duttonii* and *recurrentis*) with a high percentage of identity (100%) and a bootstrap value of 71%. In West Africa, *B. crocidurae* causes severe borreliosis, a rodent-borne disease transmitted by ticks and lice [52]. The *Ehrlichia* sequences were 100% identical to and clustered with the recently described Candidatus *Ehrlichia khabarensis* isolated from voles and shrews in the Far East of Russia [53]. The *Rickettsia* sequences were 100% identical to the sequence of *R. typhi*, a species of the typhus group responsible for murine typhus [54], but this clade was only weakly differentiated from many other *Rickettsia* species (bootstrap support of 61%). The most likely species corresponding to the sequences of the *Streptobacillus* OTU was S. *moniliformis*, with a high percentage of identity (100%) and a bootstrap value of 100%. This bacterium is common in rats and mice and causes a form of rat-bite fever, Haverhill fever [55]. The *Orientia* sequences corresponded to *O. chuto*, with a high percentage of identity (100%) and a bootstrap value of 77%. This species was recently isolated from a patient infected in Dubai [56]. Finally, accurate species determination was not possible for *Bartonella*, as the 16S rRNA gene does not resolve the species of this genus well [57]. Indeed, the sequences from the *Bartonella* OTU detected in our rodents corresponded to at least seven different species *(elizabethae, japonica, pachyuromydis, queenslandis, rattaustraliani, tribocorum, vinsonii)* and a putative new species recently identified in Senegalese rodents [58].

**Table 3.**
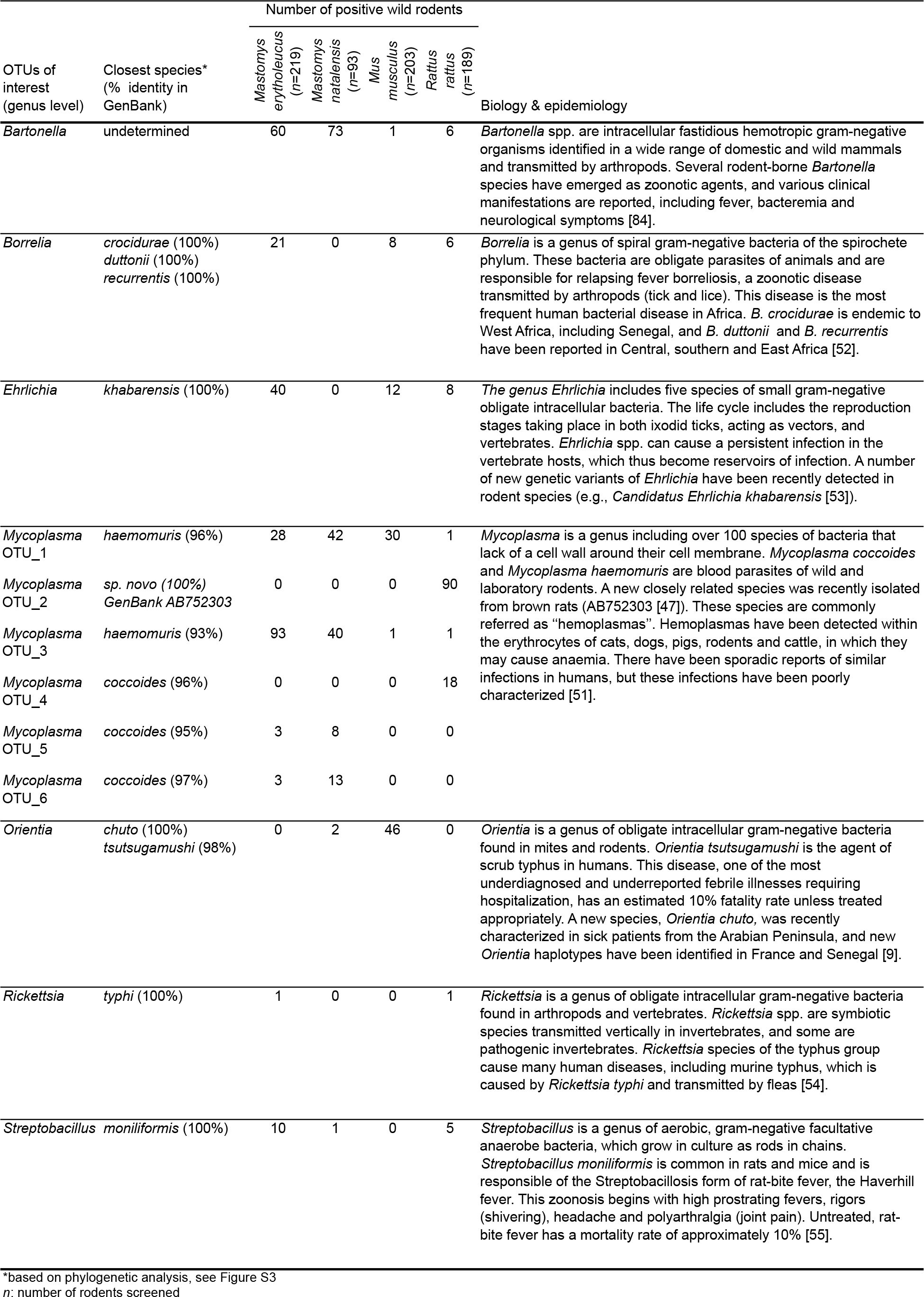
**Detection of 12 bacterial OTUs in the four wild rodent species (n=704) sampled in Senegal;** biology and pathogenicity of the corresponding bacterial genus. *n=*number of rodents analyzed.

These findings demonstrate the considerable potential of 16S rRNA amplicon sequencing for the rapid identification of zoonotic agents in wildlife, provided that the post-sequencing data are cleaned beforehand. *Borrelia* [52] and *Bartonella* [58] were the only ones of the seven pathogenic bacterial genera detected here in Senegalese rodents to have been reported as present in rodents from West Africa before. The other bacterial genera identified here have previously been reported to be presented in rodents only in other parts of Africa or on other continents. *Streptobacillus moniliformis* has recently been detected in rodents from South Africa [59] and there have been a few reports of human streptobacillosis in Kenya [60] and Nigeria [61]. *R. typhi* was recently detected in rats from Congo, in Central Africa [62], and human seropositivity for this bacterium has been reported in coastal regions of West Africa [63]. With the exception of one report in Egypt some time ago [64], *Mycoplasma* has never before been reported in African rodents. Several species of *Ehrlichia* (from the *E. canis* group: *E. chaffeensis, E. ruminantium, E. muris, E. ewingii*) have been characterized in West Africa, but only in ticks from cattle [65] together with previous reports of possible cases of human ehrlichioses in this region [66]. Finally, this study reports the first identification of *Orientia* in African rodents [9]. There have already been a few reports of suspected human infection with this bacterium in Congo, Cameroon, Kenya and Tanzania [67].

***Estimating prevalence and coinfection.*** After data filtering, we were able to estimate the prevalence in rodent populations and to assess coinfection in individual rodents, for the 12 bacterial OTUs. Bacterial prevalence varied considerably between rodent species (Table 3). *Bartonella* was highly prevalent in the two multimammate rats *M. natalensis* (79%) and *M. erythroleucus* (27%); *Orientia* was prevalent in the house mouse *M. musculus* (22%) and *Ehrlichia* occurred frequently in only one on the two multimammate rats *M. erythroleucus* (18%). By contrast, the prevalence of *Streptobacillus* and *Rickettsia* was low in all rodent species (<5%). Coinfection was common, as 184 rodents (26%) were found to be coinfected with bacteria from two (19%), three (5%), four (2%) or five (0.1%) different bacterial pathogens.

**Figure 5.**
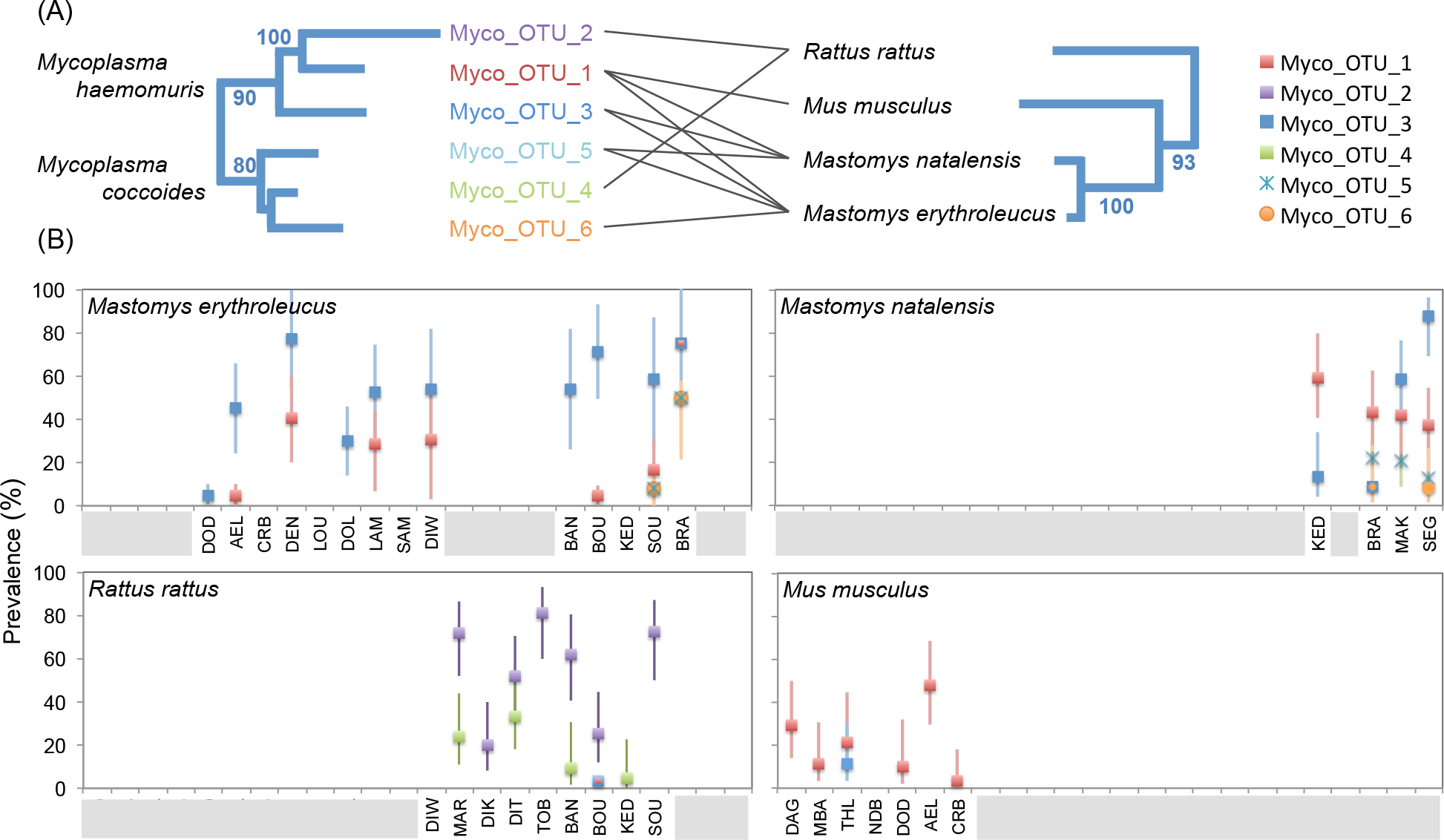
**Prevalence of *Mycoplasma* lineages in Senegalese rodents, by site, and 427 phylogenetic associations between *Mycoplasma* lineages and rodent species.** (A)Comparison of phylogenetic trees based on the 16S rRNA V4-sequences of *Mycoplasma*, and on the mitochondrial cytochrome *b* gene and the two nuclear gene fragments (IRBP exon 1 and GHR) for rodents (rodent tree redrawn from [93]). Lines link the *Mycoplasma* lineages detected in the various rodent species (for a minimum site prevalence exceeding 10%). The numbers next to branches are bootstrap values (only shown if >70%). (B) Plots of OTU prevalence with 95% confidence intervals calculated by Sterne's exact method [94] by rodent species and site (see [69] for more information about site codes and their geographic locations). The gray bars in the X-legend indicate sites from which the rodent species concerned is absent.

Interestingly, several *Mycoplasma* OTUs appeared to be specific to a rodent genus or species (Table 3; Figure 5, Panel A). OTU_2, putatively identified as a recently described lineage isolated from brown rat, *Rattus norvegicus* [47], was specifically associated with *R. rattus* in this study. Of the OTUs related to *M. coccoides*, OTU_4 was found exclusively in *R. rattus*, whereas OTUs_5 and 6 seemed to be specific to the two multimammate rats (*M. erytholeucus and M. natalensis)*. Comparative phylogenies of *Mycoplasma* OTUs and rodents showed that *R. rattus*, which is phylogenetically more distantly related to the other three rodents, contained a *Mycoplasma* community different from that in the *Mus-Mastomys* rodent clade (Figure 5, Panel A). Pathogen prevalence also varied considerably between sites, as shown for the six *Mycoplasma* OTUs (Figure 5, Panel B). This suggests that the infection risks for animals and humans vary greatly according to environmental characteristics and/or biotic features potentially related to recent changes in the distribution of rodent species in Senegal [68,69].

## Perspectives

***Recommendation for future experiments.*** Our experiments demonstrated the need to include many different kind of controls, at different steps, in order to avoid data misinterpretation. In particular, alien positive controls are important for establishing threshold values for OTUs positivity. These alien positive controls should include taxa distant enough from potential pathogens in order to avoid potential confusion between sequences of alien controls and sequences that result from actual infection of rodent samples. Ideally, one should choose alien positive controls from bacterial genera which are not able to infect the study’;s host species. In our study, the use of *Mycoplasma* and *Borrelia* species as alien positive controls was not ideal because both genera are potential rodent pathogens. Thankfully, the species used as alien controls could be easily distinguished from the species found in rodents on the basis of the phylogenetic analyses of the V4 sequences. However, based on our experience, we recommend using bacterial genera phylogenetically distant from pathogenic genera as alien controls when possible.

The inclusion of negative controls of DNA extraction in studies based on massive sequencing of 16S rRNA amplicons had long been overlooked, until the publication of Salter in 2014 [23] demonstrated the high pollution of laboratory reagents by bacterial DNA. Most studies published prior to this reported no extraction controls in their protocols. Here, we have performed one negative control for extraction per DNA extraction microplate; with each run consisting of four DNA extraction microplates, and each control having been analyzed in two replicate, we have a total of 8 negative controls for extraction per run which are analyzed twice. Based on our experience, we recommend performing at least this number of extraction controls per run. Further increases in the number of extraction controls per microplate would further improve the efficiency of data filtering and so the quality of the data produced.

The protocol of PCR amplification is also of importance for insuring data quality. In our study, we built separate amplicon libraries for each sample separately, and used very long PCR elongation times (5min) in order to mitigate the formation of chimeric reads [18] (also called jumping PCR). High numbers of PCR cycles are also known to increase chimera formation, yet as mentioned by Schnell et al [70], this parameter is mainly only critical when bulk amplification of pools of tagged/indexed amplicons is performed (e.g. when using the Illumina TrueSeq library preparation kit). As we used separate amplicon libraries for each sample, we believe that the relatively high number of PCR cycles we used (40 cycles) had minimal impact on chimera formation, and this protocol ensures the absence of chimeric sequences between samples. We had chosen to maximize the number of cycles to enhance our ability to detect pathogenic bacteria, which are sometimes in low quantity in animal samples. Fine-tuning the balance between these parameters deserves further study.

Moreover, in our study we targeted the spleen to detect bacterial infections based on the fact that this organ is known to filter microbial cells in mammals. However we lack the data to be certain that the spleen is the best organ for giving a global picture of bacterial infection in rodents (and more broadly, vertebrates). We are currently conducting new experiments to address this issue.

Finally, in our experiments, about a third of OTU sequences were attributed neither to contamination nor to (known) pathogenic genera. We currently have no precise idea of the significance of the presence of these OTUs in the rodent spleens. Part of these OTUs could be linked to further undetected biases during data generation; in spite of all the precautions we have implemented here, other biases may still elude detection. Such biases could explain the very high numbers of rare OTUs (11,947 OTUs < 100 reads), which together represent more than 88% of the total number of OTUs but less than 1 % of the total number of sequences (both runs combined).

Additionally, the presence of an OTU in a rodent spleen does not necessarily imply that the OTU is pathogenic. We know little about the microbiome of healthy vertebrates organs, yet the sharp increase of microbiome studies over the last few years has led to the discovery that microbiota communities appear to be specific to each part of the vertebrate’s body, including internal tissues and blood [71] The OTUs detected in rodent’s spleen could thus simply be part of the healthy microbiome of the organ. These issues deserve better documentation. Our results thus pave the way for future research on unknown bacterial pathogens and the microbiome of healthy organs in vertebrates.

***Improving HTS for epidemiological surveillance.*** The screening strategy described here has the considerable advantage of being non-specific, making it possible to detect unanticipated or novel bacteria. Razzauti *et al*. [8] recently showed that the sensitivity of 16S rRNA amplicon sequencing on the MiSeq platform was equivalent to that of whole RNA sequencing (RNAseq) on the HiSeq platform for detecting bacteria in rodent samples. However, little is known about the comparative sensitivity of HTS approaches relative to qPCR with specific primers, the current gold standard for bacterial detection within biological samples. Additional studies are required to address this question. Moreover, as 16S rRNA amplicon sequencing is based on a short sequence, it does not yield a high enough resolution to distinguish between species in some bacterial genera, such as *Bartonella*, nor to distinguishing between pathogenic and non-pathogenic strains within the same bacterial species.

To get this information, we thus need to follow up the 16S rRNA amplicon sequencing with complementary laboratory work. Whole-genome shotgun or RNAseq techniques provide longer sequences, through the production of longer reads or the assembly of contigs, and they might therefore increase the accuracy of species detection [72]. However, these techniques would be harder to adapt for the extensive multiplexing of samples [8]. Other methods could be used to assign sequences to bacterial species or strains for samples found positive for a bacterial genus following the 16S rRNA screening. For example, positive PCR assays could be carried out with bacterial genus-specific primers, followed by amplicon sequencing, as commonly used in MLSA (multilocus sequence analysis) strategies [73] or high-throughput microfluidic qPCR assays based on bacterial species-specific primers could be used [74]. High-throughput amplicon sequencing approaches could be fine-tuned to amplify several genes for species-level assignment, such as the *gltA* gene used by Gutierrez *et al*. [75] for the *Bartonella* genus, in parallel with the 16S rRNA-V4 region.

This strategy could also easily be adapted for other microbes, such as protists, fungi and even viruses, provided that universal primers are available for their detection (see [76,77] for protists and fungi, and [78] for degenerate virus family-level primers for viruses). Finally, our filtering method could also be translated to any other postsequencing dataset of indexed or tagged amplicons in the framework of environmental studies (e.g. metabarcoding for diet analysis and biodiversity monitoring [79], the detection of rare somatic mutations [80] or the genotyping of highly polymorphic genes (e.g. MHC or HLA typing, [81,82]).

***Monitoring the risk of zoonotic diseases.*** Highly successful synanthropic wildlife species, such as the rodents studied here, will probably play an increasingly important role in the transmission of zoonotic diseases [83]. Many rodent-borne pathogens cause only mild or undifferentiated disease in healthy people, and these illnesses are often misdiagnosed and underreported [55,84–87]. The information about pathogen circulation and transmission risks in West Africa provided by this study is important in terms of human health policy. We show that rodents carry seven major pathogenic bacterial genera: *Borrelia, Bartonella, Mycoplasma, Ehrlichia, Rickettsia, Streptobacillus* and *Orientia*. The last five of these genera have never before been reported in West African rodents. The data generated with our HTS approach could also be used to assess zoonotic risks and to formulate appropriate public health strategies involving the focusing of continued pathogen surveillance and disease monitoring programs on specific geographic areas or rodent species likely to be involved in zoonotic pathogen circulation, for example.

## Materials & Methods

***Ethics statement***. Animals were treated in accordance with European Union guidelines and legislation (Directive 86/609/EEC). The CBGP laboratory received approval (no. B 34–169–003) from the Departmental Direction of Population Protection (DDPP, Hérault, France), for the sampling of rodents and the storage and use of their tissues. None of the rodent species investigated in this study has protected status (see UICN and CITES lists).

***Sample collection***. We sampled rodents in 24 villages of the Sahelian and Sudanian climatic and biogeographical zones in Senegal (see Dalecky et al. [69] for details on the geographic location and other information on the villages). Rodents were sampled by live trapping according to the standardised protocol described by Dalecky et al. [69]. Briefly, traps were set within homes (one single-capture wire-mesh trap and one Sherman folding box trap per room) during one to five consecutive days. Each captured rodent was collected alive and transported to the field laboratory. There, rodents were killed by cervical dislocation, as recommended by Mills *et al*. [88] and dissected as described in Herbreteau *et al*. [89]. Rodent species were identified by morphological and/or molecular techniques [69]. The information concerning the rodent collection (sample ID, locality and species) is provided in the Table S2. Cross-contamination during dissection was prevented by washing the tools used successively in bleach, water and alcohol between rodents. We used the spleen for bacterial detection, because this organ is a crucial site of early exposure to bacteria [90]. Spleens were placed in RNAlater (Sigma) and stored at 4°C for 24 hours and then at −20°C until their use for genetic analyses.

***Target DNA region and primer design***. We used primers with sequences slightly modified from those of the universal primers of Kozich *et al*. [18] to amplify a 251-bp portion of the V4 region of the 16S rRNA gene (16S-V4F: GTGCCAGCMGCCGCGGTAA; 16S-V4R: GGACTACHVGGGTWTCTAATCC). The ability of these primers to hybridize to the DNA of bacterial zoonotic pathogens was assessed by checking that there were low numbers of mismatched bases over an alignment of 41,113 sequences from 79 zoonotic genera inventoried by Taylor et al [1], extracted from the Silva SSU database v119 [44]. The FASTA file is available in the Dryad Digital Repository http://dx.doi.org/10.5061/dryad.m3p7d [42].

We used a slightly modified version of the dual-index method of Kozich *et al*. [18] to multiplex our samples. The V4 primers included different 8-bp indices (called i5 index in the forward and i7 index in the reverse) and Illumina adapters (called P5 adapter in the forward and P7 adapter in the reverse) in the 5’ position. The combinations of 24 i5-indexed primers and 36 i7-indexed primers made it possible to identify 864 different PCR products loaded onto the same MiSeq flowcell. Each index sequence differed from the others by at least two nucleotides, and each nucleotide position in the sets of indices contained approximately 25% of each base, to prevent problems due to Illumina low-diversity libraries (Table 1).

***DNA extraction and PCRs***. All pre-PCR laboratory manipulations were conducted with filter tips under a sterile hood in a DNA-free room, i.e. room dedicated to the preparation of PCR mix and equipped with hoods that are kept free of DNA by UV irradiation and bleach treatment. DNA from bacterial isolates (corresponding to DNA extracts from laboratory isolates of *Bartonella taylorii, Borrelia burgdorferi* and *Mycoplasma mycoides)* was extracted in another laboratory, and PCRs from these isolates were performed after the amplifications of the DNA from rodents to avoid cross-contamination between samples and bacterial isolates. DNA was extracted with the DNeasy 96 Tissue Kit (Qiagen) with final elution in 200 μl of elution buffer. One extraction blank (NC_ext_), corresponding to an extraction without sample tissue, was systematically added to each of the eight DNA extraction microplates. DNA was quantified with a NanoDrop 8000 spectrophotometer (Thermo Scientific), to confirm the presence of a minimum of 10 ng/μl of DNA in each sample. DNA amplification was performed in 5 μl of Multiplex PCR Kit (Qiagen) Master Mix, with 4 of combined i5 and i7 primers (3.5μM) and 2 μl of genomic DNA. PCR began with an initial denaturation at 95°C for 15 minutes, followed by 40 cycles of denaturation at 95°C for 20 s, annealing at 55°C for 15 s and extension at 72°C for 5 minutes, followed by a final extension step at 72°C for 10 minutes. PCR products (3 μl) were verified by electrophoresis in a 1.5% agarose gel. One PCR blank (NC_pcr_), corresponding to the PCR mix with no DNA, was systematically added to each of the 18 PCR microplates. DNA was amplified in replicate for all wild rodent samples (n=711) (summary Table S1 and details by sample Table S2).

***Library preparation and MiSeq sequencing***. Two Illumina MiSeq runs were conducted. Run 1 included the PCR products (two or three replicates per sample) from wild rodents collected in north Senegal (148 *Mastomys erythroleucus* and 207 *Mus musculus*) plus the positive controls and the negative controls. Run 2 included the PCR products (two replicates per samples) from wild rodents collected in south Senegal (73 *Mastomys erythroleucus*, 93 *Mastomys natalensis* and 190 *Rattus rattus)* plus the positive controls and the negative controls. Full details on the composition of runs are given in Table S2. The MiSeq platform was chosen because it generates lower error rates than other HTS platforms [91]. The number of PCR products multiplexed was 823 for the first MiSeq run and 746 for the second MiSeq run (Table S2). Additional PCR products from other projects were added to give a total of 864 PCR products per run. PCR products were pooled by volume for each 96-well PCR microplate: 4 μl for rodents and controls, and 1.5 μl for bacterial isolates. Mixes were checked by electrophoresis on 1.5% agarose gels before their use to generate a “super-pool” of 864 PCR products for each MiSeq run. We subjected 100 μl of each “super-pool” to size selection for the full-length amplicon (V4 hypervariable region expected median size: 375 bp including primers, indexes and adaptors and 251bp excluding primers, indexes and adaptors), by excision from a low-melting agarose gel (1.25%) to discard non-specific amplicons and primer dimers. A PCR Clean-up Gel Extraction kit (Macherey-Nagel) was used to purify the excised bands. DNA was quantified by using the KAPA library quantification kit (KAPA Biosystems) on the final library before loading on a MiSeq (Illumina) flow cell (expected cluster density: 700-800 K/mm^2^) with a 500-cycle Reagent Kit v2 (Illumina). We performed runs of 2’ 251 bp paired-end sequencing, which yielded high-quality sequencing through the reading of each nucleotide of the V4 fragments twice after the assembly of reads 1 and reads 2. The raw sequence reads (.fastq format) are available in the Dryad Digital Repository http://dx.doi.org/10.5061/dryad.m3p7d [42].

***Bioinformatic and taxonomic classification.*** MiSeq datasets were processed with mothur v1.34 [43] and with the MiSeq standard operating procedure (SOP) [18]. Briefly, the MiSeq SOP (http://www.mothur.org/wiki/MiSeqSOP) allowed us to: 1) construct contigs of paired-end read 1 and read 2 using the make.contig command; 2) remove the reads with poor quality of assembly (> 275 bp); 3) align unique sequences on the SILVA SSU Reference alignment v119 [44]; 4) remove the misaligned, non-specific (eukaryotic) and chimeric reads (uchime program); 5) regroup the reads into Operational Taxonomic Units (OTUs) with a 3% divergence threshold; and 6) classify the OTUs using the Bayesian classifier included in mothur (bootstrap cutoff = 80%) and the Silva taxonomic file. At the end of the process, we obtained a table giving the number of reads for each OTU in line and each PCR product in column. For each OTU, the taxonomic classification (up to genus level) was provided. The abundance table generated by mothur for each PCR product and each OTU was filtered as described in the Results section. The most abundant sequence for each OTU in each sample was extracted from the sequence dataset with a custom-written Perl script (available in the Dryad Digital Repository http://dx.doi.org/10.5061/dryad.m3p7d [42]). The most abundant sequences for the 12 OTUs are available from GenBank (Accession Number KU697337 to KU697350). The sequences were aligned with reference sequences from bacteria of the same genus available from the SILVA SSU Ref NR database v119, using SeaView v4 [92]. We used a neighbor-joining method (bioNJ) to produce phylogenetic trees with a Kimura 2-Parameter model using SeaView software, and species were identified on the basis of the “closest phylogenetic species”. We also used our sequences for blast analyses of GenBank (blastn against nucleotide collection (nr/nt) performed in january 2016) to identify the reference sequences to which they displayed the highest percentage identity. The raw abundance table, the mothur command lines, the mothur output files, the Perl script and the FASTA files used for the phylogenetic analyses are available in the Dryad Digital Repository http://dx.doi.org/10.5061/dryad.m3p7d [42].

## Acknowledgments

This study was funded by the French National Institute for Agricultural Research (INRA) Meta-omics and microbial ecosystems metaprogram (Patho-ID project: Rodent and tick pathobiomes), the ANR ENEMI (ANR-11-JSV7-0006) and supported by the COST Action TD1303 (EurNegVec). We would like to thank Virginie Dupuy for extracting DNA from bacterial cultures as well as Julie Sappa from Alex Edelman & Associates, Jessie L Abbate and Petra Villette for improving the English writing. Analyses were performed on the CBGP HPC computational platform. The funders had no role in study design, data collection and analysis, the decision to publish, or preparation of the manuscript.

## Authors’ contributions

The study was conceived and designed by MG and JFC. MG, AL, CT, LT, HV and MR carried out the molecular biology procedures and validated the MiSeq data. MG, EB, MB and ADG contributed to the development of bioinformatics methods and validated taxonomic assignments. JFC and MTV coordinated the Patho-ID project and CB and NC coordinated the ENEMI project. MG, JFC, LT, CB and NC analyzed the data. MG and JFC wrote the manuscript. CB, NC, MR and MVT helped to draft and to improve the manuscript. All the authors have read and approved the final manuscript.

## Supplemental material

**Figure S1.**
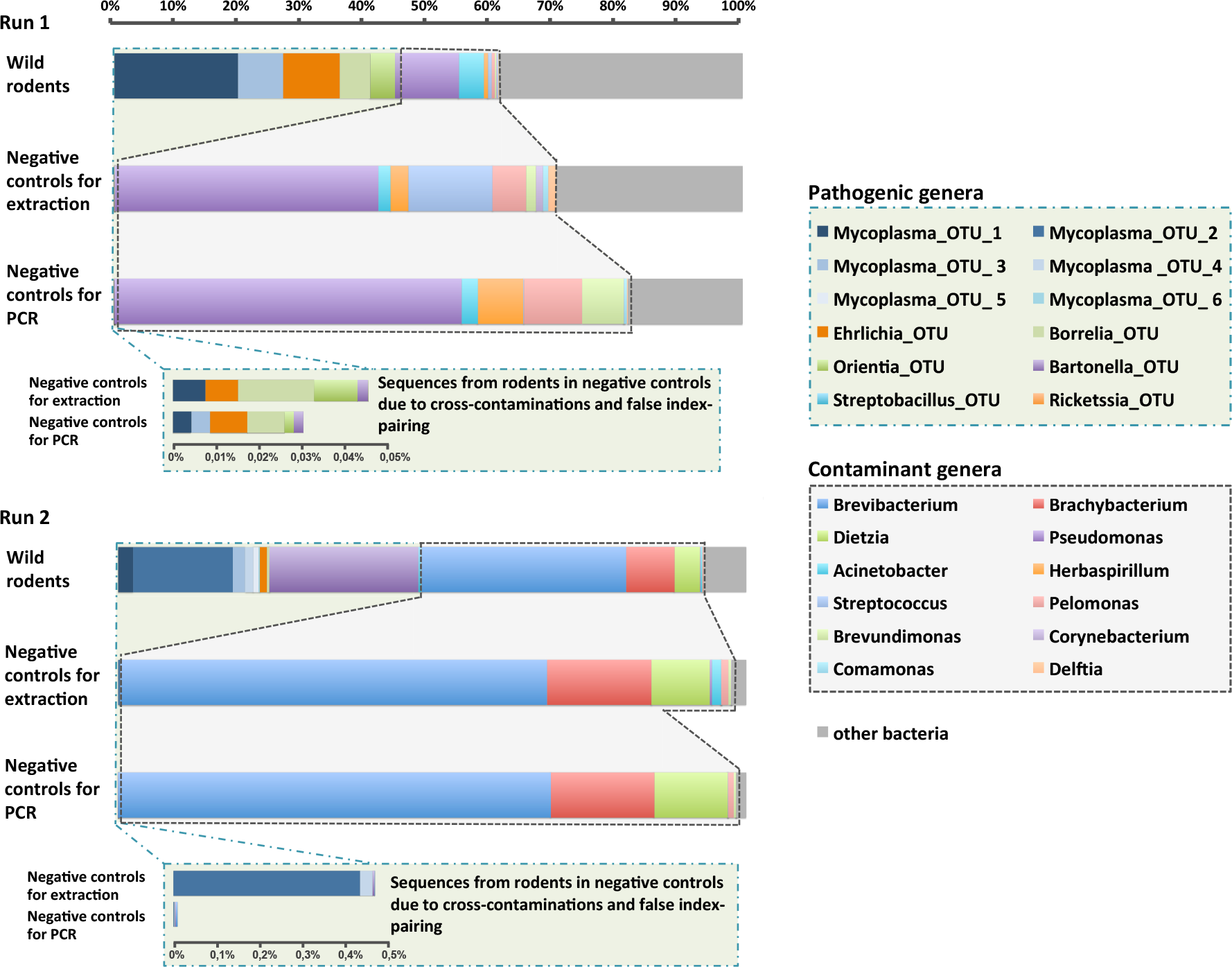
**Taxonomic assignment of the V4 16S rRNA sequences in wild rodents and in negative controls for extraction and of PCR.** The histograms show the percentage of sequences for the most abundant bacterial genera in the MiSeq run 1 and run 2. Notice the presence of several bacterial genera in the controls, which were likely due to the inherent contamination of laboratory reagents by bacterial DNA and which are thereafter called contaminant genera. These contaminant genera are also present (in lower percentage) in the rodent samples. The different in bacterial contaminant composition between run 1 and run 2 reflects the use of different kits manufactured at several months apart (Qiagen technical service, pers. com.). The differences in the pathogenic bacteria proportions and compositions between run 1 and run 2 reflects the different origins of the samples (A) run 1: *Mastomys erythroleucus* (n=148) and *Mus musculus* (n=207) from the north Senegal; (B) run 2: *Mastomys erythroleucus* (n=73), *Mastomys natalensis* (n=93) et *Rattus rattus* (n=190) from the south Senegal).

**Figure S2.**
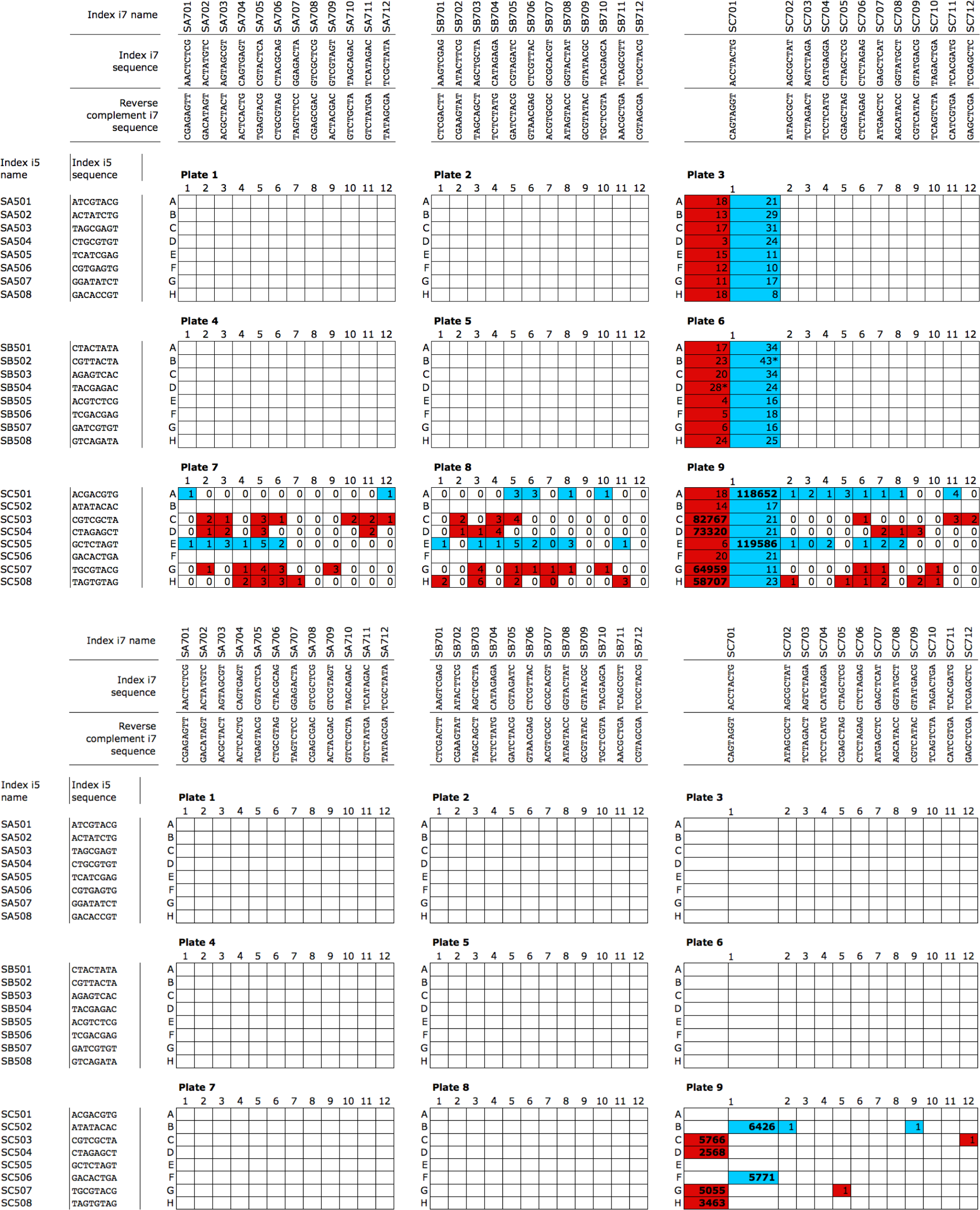
**Numbers of sequences of the positive controls for indexing PC_Borreiia_b_ (in blue) and PC_Mycoplasma_m_ (in red) in the various PCR products, with a dual-indexing design, for MiSeq runs 1 (a) and 2 (b).** The twa PCRs far PC_Barreir_b_ were performed with 96-well mieraplrte 9, pasitians A1 rnd E1 far run 1 rnd B1 rnd F1 far run 2, rnd the faur PCRs far PC_Myeaplrsmr_m_ were perfarmed with 96-well mieraplrte 9, pasitians C1, D1, G1 rnd H1 far the twa runs. The numbers af sequences far the ather wells earrespand ta indexing mistakes due ta frlse index-priring due ta mixed clusters during the sequencing (see Table 1).

**Figure S3.**
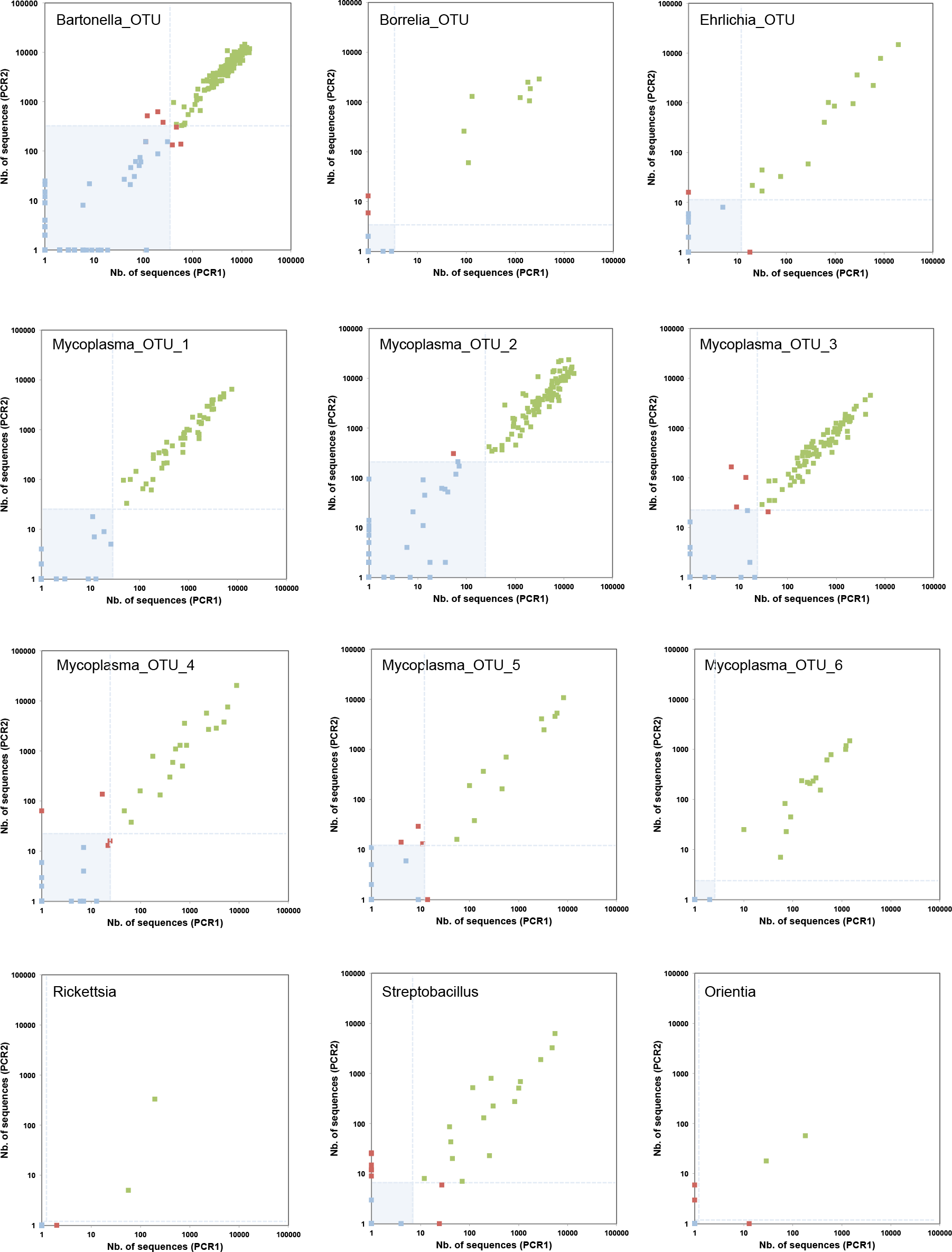
**Plots of the number of sequences (log (x+1) scale) from bacterial OTUs in both PCR replicates (PCR1 & PCR2) for the 356 wild rodents analyzed in the second MiSeq run.** Note that each rodent was tested with two replicate PCRs. Green points correspond to rodents with two positive results after the filtering process; red points correspond to rodents with one positive result and one negative result; and blue points correspond to rodents with two negative results. The light blue area and lines correspond to the threshold values used for the data filtering: samples below the lines are filtered out. See Figure 4 for plots corresponding to the first MiSeq run.

**Figure S4.**
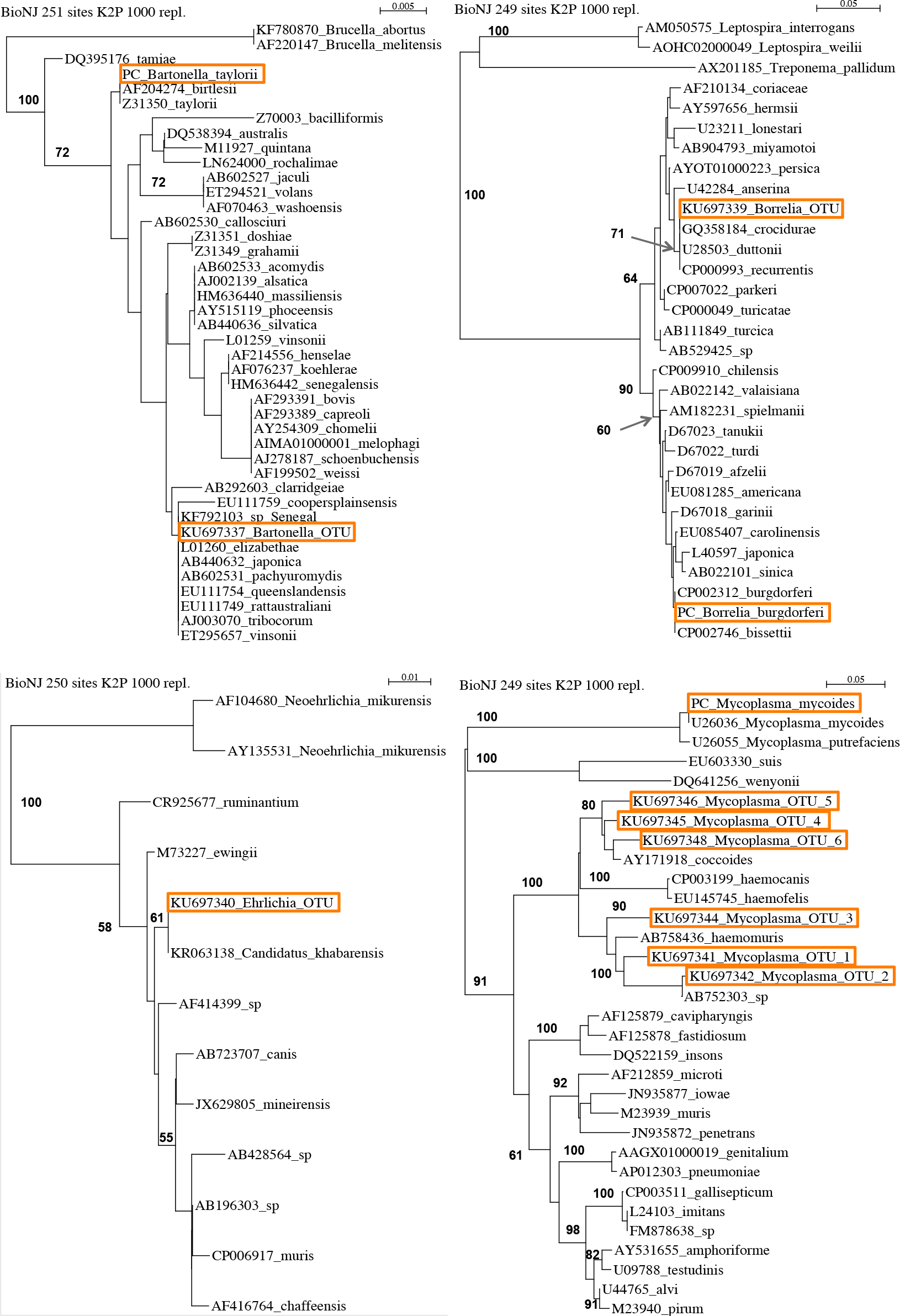
**Phylogenetic trees of the 16S rRNA V4 sequences for 12 pathogenic bacterial OTUs detected in wild rodents from Senegal.** Sequences boxed with an orange line were retrieved from African rodents and/or corresponds to positive controls (PC) for *Borellia burgdorferi, Mycoplasma mycoides* and *Bartonella taylorii*. The other sequences were extracted from the SILVA database and GenBank. Trees include all lineages collected for *Rickettsia, Bartonella, Ehrlichia* and *Orientia*, but only lineages of the Spotted Fever Group for *Borrelia*, and lineages of the pneumonia group for *Mycoplasma*. The numbers indicated are the bootstrap values >55%. Fasta files used have been deposited in the Dryad Digital Repository: http://dx.doi.org/10.5061/dryad.m3p7d.

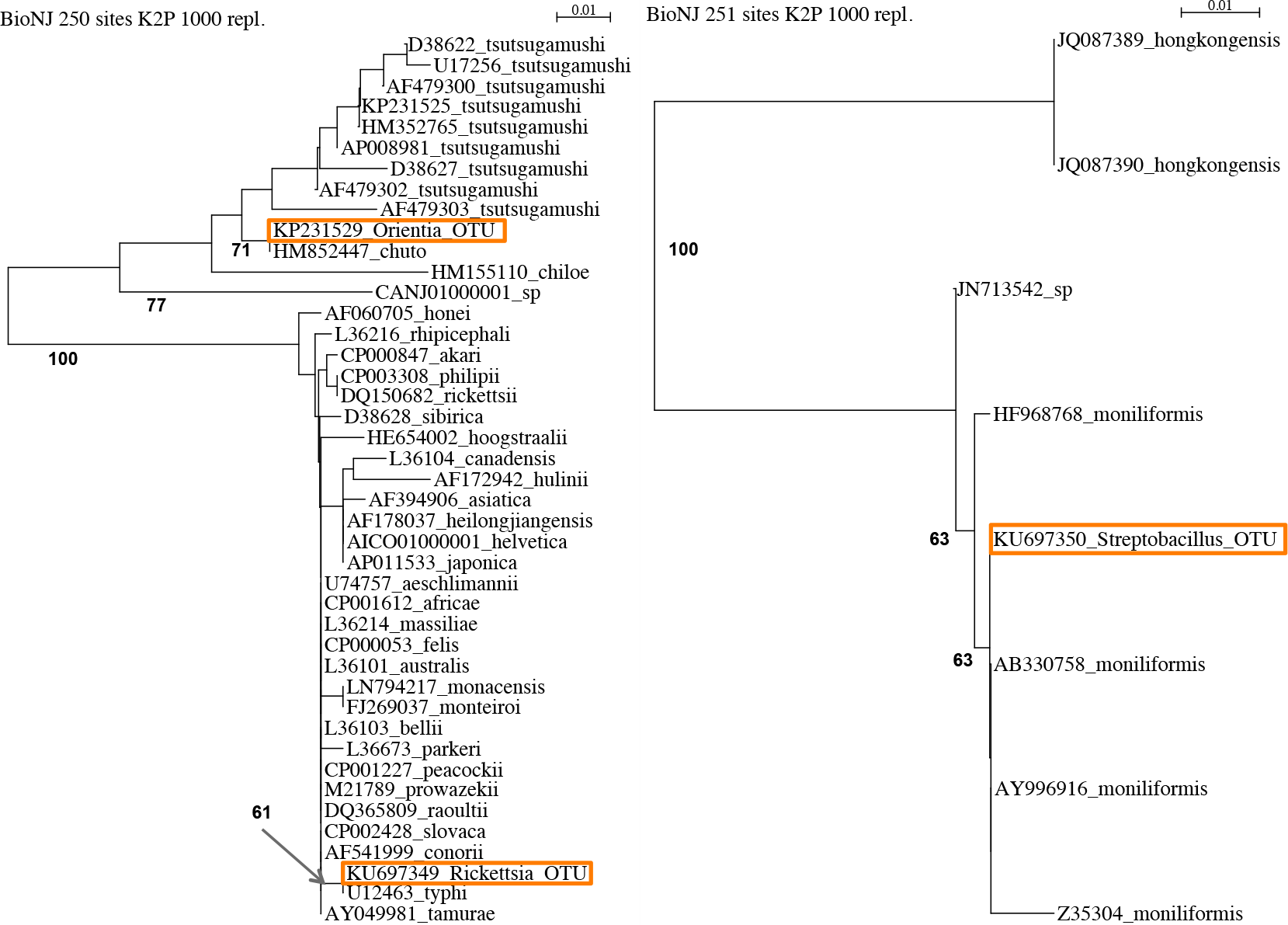

**Table S3.**
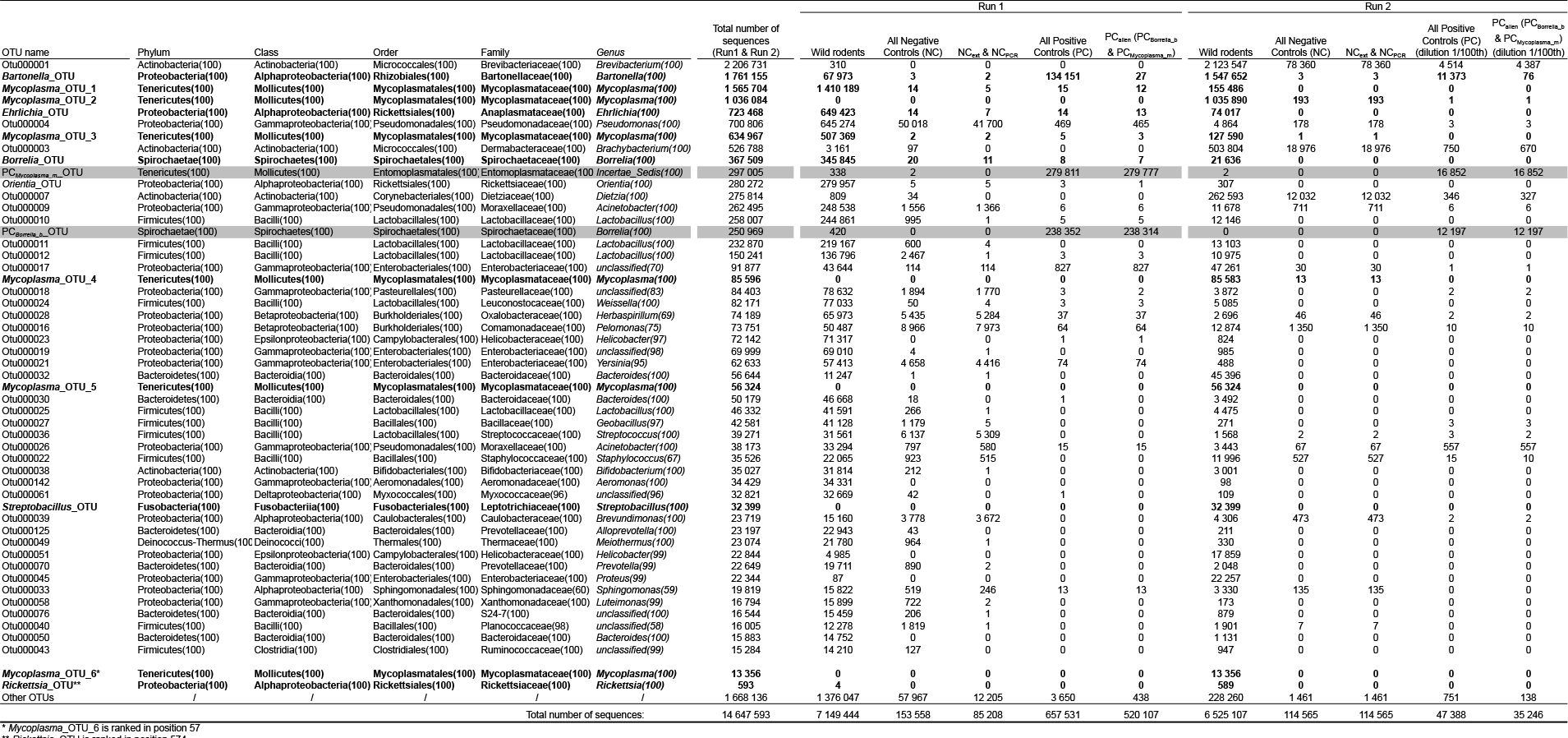
**The 50 most abundant OTUs in wild rodents and controls.** The twelve pathogenic OTUs from wild rodents are in bold and italic. The two OTUs from PCaiien (PC*Bomiia_b & PCMycopiasma_m*) are highlighted in grey. A blank space was added at the end of the table to distinguish the first 50 most abundant OTUs and the *Mycoplasma_*OTU_6 and *Rickettsia*_OTU ranked in position 57 and 574 respectively.

**Table S1.**
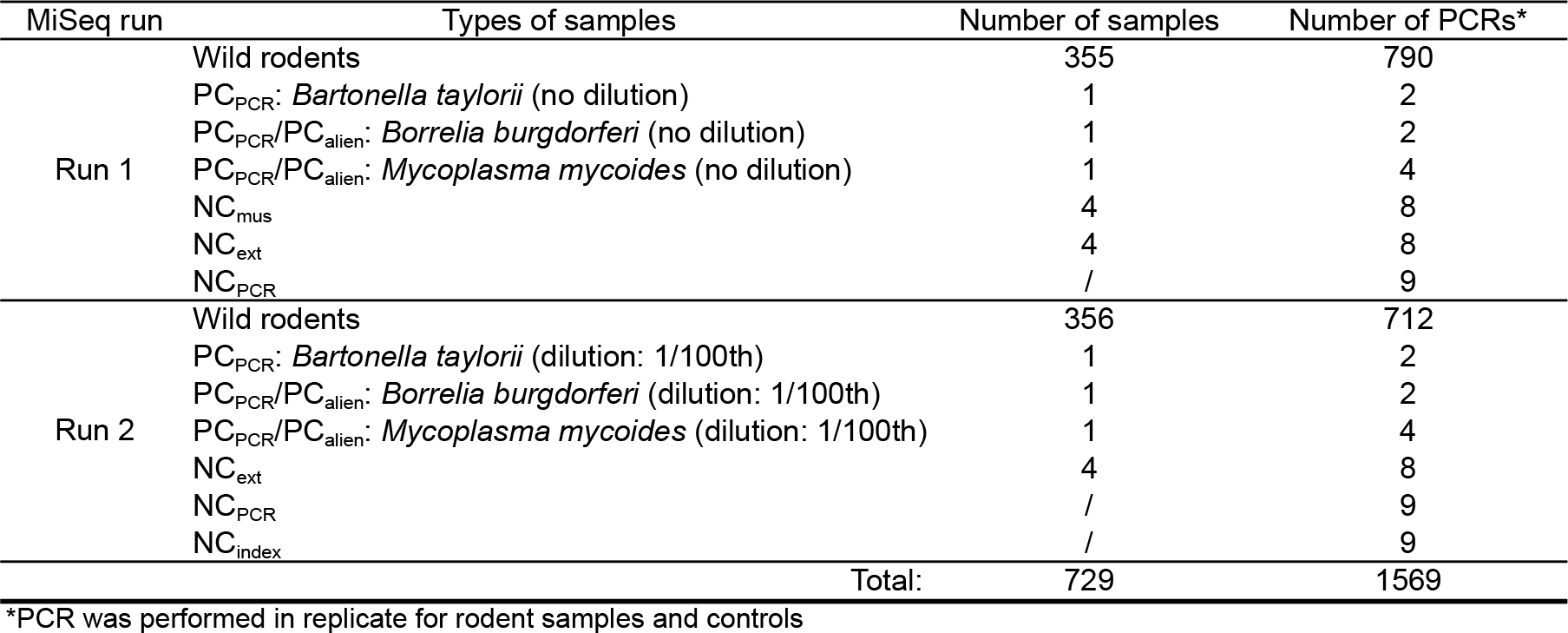
**Numbers of samples and numbers of PCRs for wild rodents and controls.** Negative Controls for dissection, NC_mus_; Negative Controls for extraction, NC_ext_; Negative Controls for PCR, NC_PCR_; Negative Controls for indexing, NC_index_; Positive Controls for PCR, PC_PCR_; Positive Controls for Indexing, PC_alien_. See also Figure 1 for more details concerning negative controls (NC) and positive controls (PC). See also Figure 1 and Box 1.

**Table S4.**
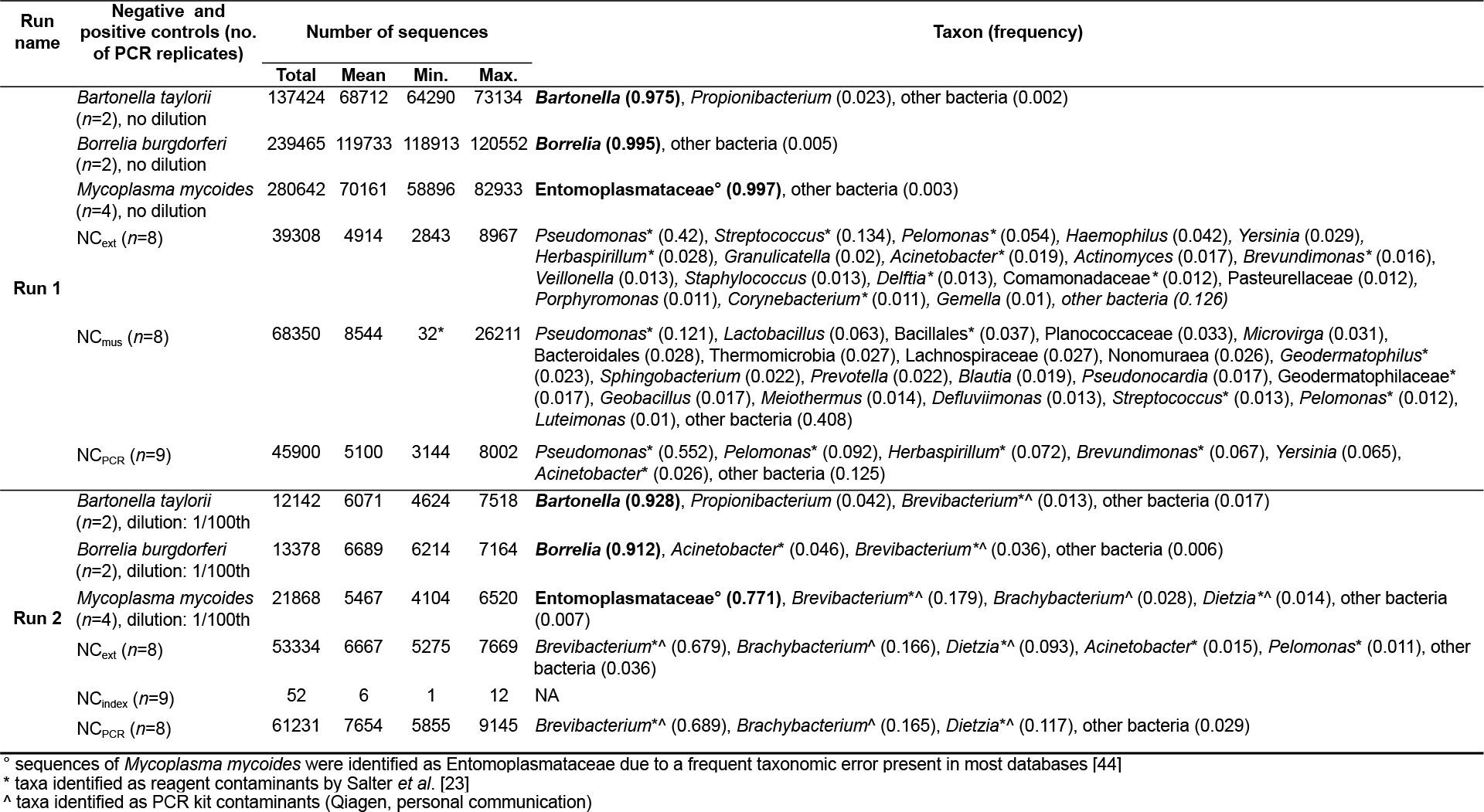
**Bacterial contaminants observed in negative ad positive controls.** They were identified as contaminants on the basis of negative controls for extraction and PCR. Taxa in bold correspond to the sequences of DNA extracted from laboratory isolates.

**Table S5.**
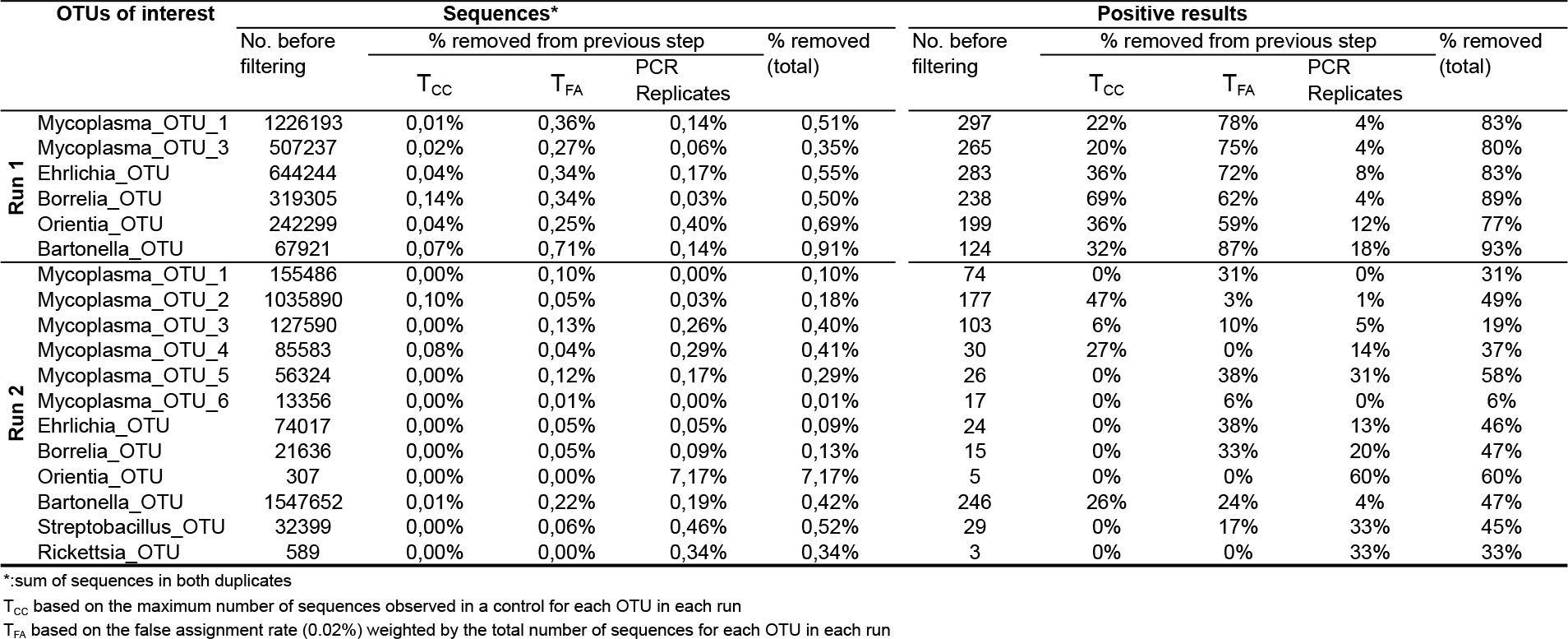
**Proportion of sequences and proportion of positive results removed at each step in data filtering.** Note that severa positive results may be recorded for the same rodent in cases of co-infection.

**Table S6.** **Proportion of positive results for both PCR products at each step in data filtering.** Note that several positive results may be recorded for the same rodent in cases of co-infection.

| OTUs of interest |  | % of rodents positive for both PCR replicates |  |  |
| --- | --- | --- | --- | --- |
|  |  | Before filtering | T<sub>CC</sub> | T<sub>FA</sub> |
| Run 1 | Mycoplasma_OTU_1 | 68% | 64% | 96% |
|  | Mycoplasma_OTU_3 | 49% | 46% | 96% |
|  | Ehrlichia_OTU | 56% | 56% | 92% |
|  | Borrelia_OTU | 38% | 53% | 96% |
|  | Orientia_OTU | 43% | 54% | 88% |
|  | Bartonella_OTU | 19% | 20% | 82% |
| Run 2 | Mycoplasma_OTU_1 | 76% | 76% | 100% |
|  | Mycoplasma_OTU_2 | 59% | 96% | 99% |
|  | Mycoplasma_OTU_3 | 86% | 92% | 95% |
|  | Mycoplasma_OTU_4 | 77% | 91% | 82% |
|  | Mycoplasma_OTU_5 | 62% | 62% | 69% |
|  | Mycoplasma_OTU_6 | 94% | 94% | 100% |
|  | Ehrlichia_OTU | 58% | 58% | 87% |
|  | Borrelia_OTU | 53% | 53% | 80% |
|  | Orientia_OTU | 40% | 40% | 40% |
|  | Bartonella_OTU | 66% | 83% | 96% |
|  | Streptobacillus_OTU | 59% | 59% | 67% |
|  | Rickettsia_OTU | 67% | 67% | 67% |

T<sub>CC</sub> based on the maximum number of sequences observed in a control for each OTU in each runT<sub>FA</sub> based on the false assignment rate (0.02%) weighted by the total number of sequences for each OTU in each run

**Table S7.** **Number of mismatches between PCR forward and reverse primers and 41,113 bacterial 16S rRNA V4 sequences of 79 zoonotic genera.** Bacterial genera were selected according to the inventory of Taylor et al [1] and sequences were extracted from the Silva SSU database v119. Numbers of mismatches > 3 correspond to sequences of bad quality from diverse taxa. The number of primer mismatches in the 10 bases of the 3′ side was < 2 for 99.93% of the reference sequences.

| Forward primer |  | Reverse primer |  |
| --- | --- | --- | --- |
| No. of mismatches | No. of sequences | No. of mismatches | No. of sequences |
| 0 | 40063 | 0 | 39901 |
| 1 | 841 | 1 | 967 |
| 2 | 101 | 2 | 132 |
| 3 | 42 | 3 | 43 |
| 4 | 8 | 4 | 24 |
| 5 | 8 | 5 | 8 |
| 6 | 6 | 6 | 4 |
| 7 | 3 | 7 | 4 |
| 8 | 2 | 8 | 4 |
| 9 | 1 | 9 | 1 |
| 10 | 4 | 10 | 1 |
| 11 | 0 | 11 | 3 |
| 12 | 0 | 12 | 1 |
| 13 | 0 | 13 | 0 |
| 14 | 0 | 14 | 1 |
| 15 | 0 | 15 | 0 |
| 16 | 0 | 16 | 0 |
| 17 | 0 | 17 | 0 |
| 18 | 0 | 18 | 0 |
| 19 | 0 | 19 | 0 |
| NA* | 34 | 20 | 0 |
|  |  | 21 | 0 |
|  |  | 22 | 0 |
|  |  | NA* | 19 |

\* Partial sequences for the primer region

